# Small and large extracellular vesicles from human preovulatory follicular fluid display distinct ncRNA cargo profile and differential effect on granulosa cell line KGN

**DOI:** 10.1101/2025.02.13.638014

**Authors:** Inge Varik, Katariina Johanna Saretok, Kristine Rosenberg, Ileana Quintero, Maija Puhka, Nataliia Volkova, Aleksander Trošin, Paolo Guazzi, Agne Velthut-Meikas

**Affiliations:** Department of Chemistry and Biotechnology, Tallinn University of Technology, Tallinn, Estonia; Nova Vita Clinic, Tallinn, Estonia; HiPREP Core, Institute for Molecular Medicine Finland (FIMM), University of Helsinki, Helsinki, Finland; East Tallinn Central Hospital, Tallinn, Estonia; HansaBioMed Life Sciences Ltd, Tallinn, Estonia; Institute for Problems of Cryobiology and Cryomedicine, National Academy of Sciences of Ukraine, Kharkiv, Ukraine

**Keywords:** ovary, follicular fluid, extracellular vesicles, granulosa cells, ncRNA, miRNA

## Abstract

Follicular fluid extracellular vesicles (FF EVs) facilitate communication between oocytes and somatic cells within the ovarian follicle, playing a pivotal role in follicular development. This study highlights the molecular and functional distinctions between small (SEV) and large (LEV) FF EV subpopulations, revealing their specialized regulatory roles in granulosa cell (GC) biology and their consequential impact on ovarian function. Single-EV profiling uncovered distinct tetraspanin distributions, with LEVs containing a lower proportion of CD9/CD63/CD81-positive particles compared to SEVs. Functionally, SEVs reduced estradiol secretion by GCs, whereas LEVs enhanced progesterone production, demonstrating their differential effects on steroidogenesis. Transcriptomic analysis revealed extensive SEV-induced changes in GC gene expression, affecting pathways involved in transcription, TGF-β signaling, extracellular matrix (ECM) remodeling, and cell cycle regulation. In contrast, LEVs elicited minimal transcriptional changes, primarily modulating genes associated with immune regulation and oxidative stress defense. Small RNA sequencing further revealed distinct non-coding RNA (ncRNA) profiles, with SEVs enriched in miRNAs targeting pathways critical for GC differentiation, while LEVs carried higher levels of piRNAs implicated in maintaining genomic stability. These findings advance our understanding of FF EV-mediated intercellular communication and underscore the importance of investigating EV subpopulations independently.

## INTRODUCTION

The primary function of the human ovary is to produce fertilization-competent oocytes and steroid hormones, such as estrogens, progesterone, and testosterone. Its functional unit, the ovarian follicle, comprises an oocyte surrounded by granulosa cells (GCs) and theca cells (Edson et al., 2009). Follicular development progresses through multiple stages, starting with primordial follicles and culminating in the formation of a preovulatory follicle, characterized by a fluid-filled antrum (Malo et al., 2024). This antrum, which delineates GCs into mural and cumulus subtypes (Khamsi & Roberge, 2001), contains follicular fluid (FF) derived from blood plasma and GC/oocyte secretions (Hennet & Combelles, 2012). FF contains various biologically active molecules, including hormones, growth factors, proteins, peptides, amino acids, and metabolites, which collectively support folliculogenesis (Revelli et al., 2009).

Effective follicle development depends on intercellular communication between somatic and germ cells (Matzuk et al., 2002). These interactions, vital for oocyte growth, maturation, and eventual release, occur via paracrine and endocrine signaling, gap junctions, and transzonal projections (Dumesic et al., 2015; Marchais et al., 2022). Previous studies have identified extracellular vesicles (EVs) in FF across multiple species, including equine, bovine, porcine, and human (da Silveira et al., 2012; Hung et al., 2015; Matsuno et al., 2017; Rodrigues et al., 2019; Rooda et al., 2020; Santonocito et al., 2014; Sohel et al., 2013), as important mediators of this communication (Marchais et al., 2022).

EVs are a heterogenous group of lipid bilayer-enclosed particles secreted by most cell types (Willms et al., 2018), serving as messengers in intercellular communication by transferring their cargo, comprising of proteins, lipids, and nucleic acids, from donor to recipient cells (Valadi et al., 2007; van Niel et al., 2018). Among their nucleic acid cargo, small non-coding RNAs (ncRNAs), such as microRNAs (miRNAs) and piwi-interacting RNAs (piRNAs), have attracted particular interest due to their roles in gene regulation (Abramowicz & Story, 2020; Liu et al., 2019). EVs are categorized, for example, into exosomes, microvesicles, and apoptotic bodies based on their biogenesis and size. Exosomes (30-150 nm), formed within multivesicular bodies (MVBs), are released when MVBs fuse with the plasma membrane (Simpson et al., 2009; van Niel et al., 2018), while microvesicles (50-1000 nm) bud directly from the plasma membrane (Muralidharan-Chari et al., 2010; van Niel et al., 2018), and apoptotic bodies (200-5000 nm) are shed from cells undergoing programmed cell death (Willms et al., 2018). EVs are often characterized by specific membrane-associated proteins, including tetraspanins CD9, CD63, and CD81 (Jankovičová et al., 2020); however, no universal EV markers are known yet (Welsh et al., 2024).

FF EVs have been implicated in supporting GC function and oocyte development. For instance, bovine FF EVs have been shown to promote GC proliferation (Hung et al., 2017), induce cumulus GC expansion (Hung et al., 2015), and protect cumulus-oocyte complexes from apoptosis and heat shock damage (Rodrigues et al., 2019). In humans, FF EV miRNAs have been linked to pathways that regulate follicle growth and oocyte maturation (Santonocito et al., 2014). Our research group has previously demonstrated that miRNA profiles in FF EVs differ significantly from those of the somatic follicular cells and bulk FF of the same follicle (Rooda et al., 2020), suggesting that EV miRNAs serve as specific means of molecular communication within the follicle. Furthermore, differences in EV miRNA cargo between patients with polycystic ovarian syndrome (PCOS) and healthy individuals (Rooda et al., 2020) highlight their potential as diagnostic biomarkers.

Despite their recognized importance, most studies have examined the collective pool of FF EVs, overlooking the diversity among EV subpopulations. Recent research has identified at least ten distinct FF EV subpopulations (Neyroud et al., 2022), each potentially fulfilling specialized roles in the ovarian follicle. For instance, small FF EV subpopulations have been shown to exhibit heterogeneous miRNA profiles (X. Wang et al., 2021), emphasizing their unique molecular characteristics. These findings underscore the need for in-depth investigations into the specific functions of FF EV subpopulations, given their potential to serve distinct functions in ovarian physiology.

However, overlapping size and density ranges between EV subpopulations, such as exosomes and microvesicles, pose challenges for their complete isolation (Brennan et al., 2020). Moreover, there are currently no molecular markers that can definitively distinguish between these two subtypes (Welsh et al., 2024). In this study, we used size exclusion chromatography (SEC) and tangential flow filtration (TFF) with a 200 nm cut-off to separate small EVs (SEVs) and large EVs (LEVs) from human FF. We hypothesize that this size separation produced SEVs that are enriched in exosomes and LEV preparations enriched in microvesicles, though some degree of cross-contamination between the subpopulations was likely.

We characterized the molecular composition, biophysical properties, and ncRNA cargo of FF EV subpopulations. Using human granulosa-like tumor cell line KGN as a model, we also assessed their functional effects on GC viability, proliferation, steroidogenesis, and gene expression (Figure 1). Our findings provide new insights into the specialized roles of FF EV subpopulations in ovarian physiology and contribute to a deeper understanding of EV-mediated intercellular communication within the ovarian follicle.

**Figure 1.**
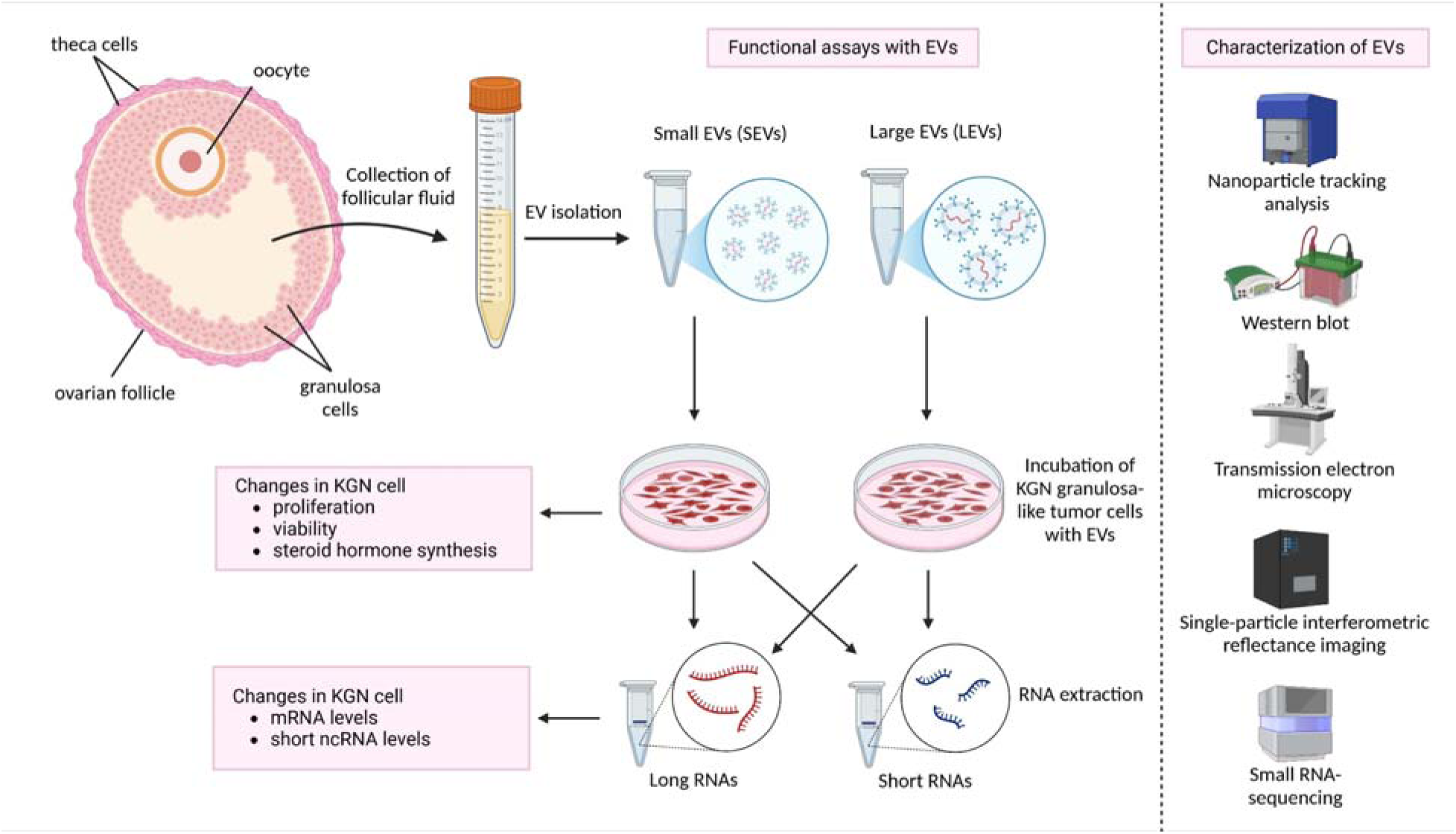
Study design. Extracellular vesicles were isolated from human ovarian follicles and separated based on size. Small and large EVs were characterized with multiple methods. Functional studies were carried out on the KGN cell line serving as a homogenous granulosa cell model. Created in BioRender. Varik, I. (2025) https://BioRender.com/p12h544.

## MATERIALS AND METHODS

### Collection of follicular fluid

FF samples were donated by patients undergoing assisted reproductive technology (ART) at Nova Vita Clinic and East Tallinn Central Hospital, both located in Tallinn, Estonia. All procedures in this study adhered to the Declaration of Helsinki and were approved by the Research Ethics Committee of the University of Tartu, Estonia (approval no. 356/M-4). Prior to participation, patients received both written and oral information about the study. Informed written consent was obtained from all subjects for the use of FF samples collected during ART treatment. To ensure confidentiality, all donated samples were anonymized before further procedures.

At Nova Vita Clinic, a gonadotropin-releasing hormone (GnRH) antagonist (Cetrotide, Merck, Darmstadt, Germany) protocol was used, along with the administration of recombinant follicle-stimulating hormone (Gonal-F, Merck; or Bemfola, Gedeon Richter Plc., Budapest, Hungary) for ovarian stimulation.

At East Tallinn Central Hospital, two ovarian stimulation protocols were employed. In long protocols, a GnRH agonist (Diphereline, Ipsen Pharma Biotech, France, Paris) was administered along with Gonal-F (Merck), Bemfola (Gedeon Richter Plc.), Ovaleap (Theramex, United Kingdom, London), Rekovelle (Ferring Pharmaceuticals, Switzerland, Saint-Prex), Pergoveris (Merck Serono), or Luveris (Merck Serono). In short protocols, a GnRH antagonist (Cetrotide, Merck) was administered, together with Gonal-F, Bemfola, Pergoveris, Ovaleap, or Rekovelle.

At both clinics, ovarian puncture was carried out transvaginally under ultrasound guidance 36 hours after human chorionic gonadotropin administration (Ovitrelle, Merck) if at least two follicles were ≥ 18 mm in diameter. The FF was collected from one or more follicles with minimal visible blood contamination and transported to the laboratory in a thermal flask. The samples were processed by centrifugation at 400 × g for 10 min to pellet intact somatic cells. The supernatant was subsequently centrifuged at 2000 × g for 10 min to remove any residual cellular debris. Finally, the cell-free FF samples were aliquoted to avoid repeated freeze-thaw cycles and stored at -80 °C until further analysis.

### Isolation of FF EVs by size exclusion chromatography

EV isolation and characterization were performed in adherence to the MISEV2023 guidelines (Welsh et al., 2024). A maxiPURE-EVs column (HansaBioMed Life Sciences, Tallinn, Estonia) was used to isolate vesicles by size exclusion chromatography (SEC). Before use, the column was washed three times with 1X Dulbecco’s phosphate-buffered saline (DPBS; Corning, Massachusetts, USA). FF samples were thawed, and pooled FF samples were prepared by combining equal volumes of FF from seven patients per pool to average out patient-to-patient variability and focus on general trends rather than individual differences between patients. In total, 42 patient samples were used to generate six pooled samples for analysis. A 20 ml aliquot of pooled FF was loaded onto the maxiPURE-EVs column and allowed to pass through by gravity flow. Fractions of 1 ml were collected sequentially, and a total of 40 fractions was obtained, using DPBS as the elution buffer. The column was washed three times with 1X DPBS between uses and reused up to five times.

### Isolation of FF EV subpopulations by tangential flow filtration

Nanoparticle tracking analysis (NTA) and total protein quantification (described in detail below) indicated that fractions 15-31 were enriched in EVs and exhibited minimal protein contamination. These fractions were pooled and subjected to ultrafiltration with a tangential flow filtration (TFF) system. A TFF-MV cartridge (HansaBioMed Life Sciences) with 200 nm pore size was employed, following the manufacturer’s instructions. Shortly, the sample was introduced into the cartridge with a clean 25 ml syringe attached to one of the TFF nozzles (syringe A). Another empty 25 ml syringe (syringe B) was connected to the second nozzle, and a collection tube was placed under the permeate nozzle. The syringes were alternatively pushed until emptied, after which syringe A was filled with 5 ml of 1X DPBS and the process was repeated. As a result, permeate containing SEVs (<200 nm) was collected. To collect the retained LEVs (>200 nm), the permeate nozzle was closed, syringe B was filled with 5 ml of 1X DPBS, re-attached to the nozzle, and both syringes were alternatively pushed 15 times. The retentate containing LEVs was collected in one of the syringes. The TFF cartridge was thoroughly washed with Milli-Q water at least three times, air-dried, and stored at room temperature (RT).

### Concentration of FF EV samples

22 ml of SEV and 5 ml of LEV samples were concentrated using 10 kDa Amicon Ultra-15 centrifugal filters (Merck Millipore, Massachusetts, USA). Samples were centrifuged at 3200 × g at RT until a final volume between 300 and 700 µl was reached. Concentrated EVs were used immediately for experiments or aliquoted and stored in premium surface tubes (04-232-3500, Nerbe Plus, Germany, Winsen) at -80 °C. The EVs were not subjected to more than one freeze-thaw cycle and were used within 18 months after storage. Upon use, EVs were thawed at RT.

### Nanoparticle tracking analysis

Size distribution and concentration of isolated EVs were measured with the ZetaView PMX 110 (Particle Metrix, Germany) instrument using its corresponding software (ZetaView 8.05.12 SP1). Before measurements, the instrument was calibrated with a 100 nm polystyrene nanoparticle standard (Applied Microspheres BV, Leusden, Netherlands) prepared in distilled water, following the manufacturer’s recommendations. Automated quality control measurements, including cell quality check and instrument alignment, were performed prior to measurements.

One milliliter of sample, diluted in 1X DPBS, was introduced into the cell, ensuring the detection of 50 to 200 particles per frame. Particle size and concentration were measured at 11 distinct positions, capturing 30 frames per position across three independent readings with a shutter speed of 100. The sensitivity for small particle measurements was set at 85, while for large particle measurements, it was set at 65. The cell temperature was maintained at 25 °C for all measurements, and the cell was washed with 1X DPBS between each measurement.

### Total protein quantification

Protein abundance was determined with the Pierce™ BCA Protein Assay Kit (Thermo Fisher Scientific, Massachusetts, USA), according to the manufacturer’s instructions, with 10 μl of undiluted intact EVs and bovine serum albumin (BSA) as a standard. The absorbance at 570 nm was measured with a TECAN GENios Pro plate reader (TECAN Group Ltd, Männedorf, Switzerland). EV purity was calculated by dividing the particle concentration (particles/ml) determined by NTA with the protein concentration (µg/ml) measured by bicinchoninic acid assay (BCA).

### Western blot

SEV and LEV samples were obtained by TFF, and fractions 32-40 (containing FF proteins) were obtained by SEC as described earlier. To precipitate proteins from 300 µl of FF, 100 µl of Milli-Q water, 400 µl of methanol (Sigma-Aldrich, Missouri, USA), and 100 µl of chloroform (Lach-Ner, Neratovice, Czech Republic) were added to the concentrated sample. The mixture was then centrifuged for 5 minutes at 14,000 g. The top layer was removed, and the protein layer was washed with 400 µl of methanol. The sample was centrifuged again as before, the protein pellet was dried, and resuspended in 0.25% SDS (SERVA Electrophoresis GmbH, Heidelberg, Germany) containing cOmplete™ Mini Protease Inhibitor Cocktail (Roche, Basel, Switzerland). For cell lysis, KGN cells were treated with RIPA buffer (50 mM Tris pH 7.4, 1% Triton X-100, 1 mM EDTA, 150 mM NaCl, 0.1% SDS) supplemented with cOmplete™ Mini Protease Inhibitor Cocktail. The lysate was maintained on ice for 30 minutes, vortexed every 10 minutes, and then centrifuged at 10,000 g for 10 minutes. Following centrifugation, the supernatant was transferred to a new tube. Lyophilized small EVs from the HCT116 cell line (HansaBioMed Life Sciences) were reconstituted in 100 μl of Milli-Q water.

10 µg of protein was mixed with either reducing 4X Laemmli buffer (250 mM Tris-HCl, 8% SDS, 40% glycerol, 0.05% bromophenol blue, 10% β-mercaptoethanol) or 4X non-reducing Laemmli buffer (250 mM Tris-HCl, 8% SDS, 40% glycerol, 0.05% bromophenol blue); the latter was used for the detection of CD9 and CD81 proteins. Samples were denatured for 5 min at 95 °C and separated by 10% SDS-PAGE at 90 V for 30 min, followed by 150 V for 1 h using the Mini-PROTEAN Tetra Vertical Electrophoresis Cell (Bio-Rad Laboratories, California, USA). Proteins were transferred onto the Immobilon®-P polyvinylidene fluoride membrane (Merck Millipore) in Towbin buffer (25 mM Tris, 192 mM glycine, 10% ethanol) at 370 mA for 1 h with the Mini Trans-Blot® cell (Bio-Rad Laboratories). The membranes were incubated in 5% non-fat dry milk in 0.1% PBST buffer (0.1% of Tween 20 in 1X DPBS) for 1 h at RT.

Immunoblotting was carried out with the following primary antibodies diluted in 2.5% non-fat dry milk in 0.1% PBST overnight at +4 °C: mouse anti-CD9 antibody (HBM-CD9-100, 1:500, HansaBioMed Life Sciences), mouse anti-CD81 antibody (HBM-CD81-EM4-100, 1:500, HansaBioMed Life Sciences), rabbit anti-HSP70 antibody (10995-1-AP-20, 1:500, Proteintech), rabbit anti-albumin antibody (16475-1-AP, 1:10,000, Proteintech, Illinois, USA), mouse anti-ApoA1 antibody (sc-376818, 1:1000, Santa Cruz Biotechnology, Texas, USA), and rabbit anti-calnexin antibody (AB2301, 1:10,000, Merck Millipore). The membranes were washed 3 × 5 min with 0.1% PBST and then incubated with either HRP-conjugated goat anti-rabbit IgG (115-035-003, 1:10,000, Jackson Immunoresearch, Pennsylvania, USA) or goat anti-mouse IgG (G-21040, 1:20,000, Thermo Fisher Scientific) diluted in 2.5% non-fat dry milk in 0.1% PBST buffer for 1 h at RT. After further washing (6 × 5 min) with 0.1% PBST and incubation with SuperSignal West ECL Substrate (Thermo Fisher Scientific) at RT, the images were acquired using the ImageQuant™ LAS 4000 instrument (GE HealthCare Technologies, Inc., Illinois, USA).

### Transmission electron microscopy

LEV (but not SEV) samples were concentrated using Amicon Ultra-0.5 Centrifugal Filter (Merck Millipore) with a 10 kDa cutoff. All EV samples were prepared for transmission electron microscopy (TEM) as in (Puhka et al., 2017). Briefly, samples with particle concentrations between 10^10^/ml and 10^11^/ml were loaded directly onto carbon-coated and glow-discharged 200 mesh copper grids with formvar or pioloform support membrane. The loaded EVs were fixed with 2% paraformaldehyde in sodium phosphate buffer (pH 7.0), washed with deionized water (PURELAB® Chorus 1 Water Purification System, ELGA LabWater), and then stained with 2% neutral uranyl acetate. Finally, the samples were embedded and further stained in 1.8% methyl cellulose (25 ctp)/0.4% uranyl acetate in deionized water on ice. The EVs were visualized by TEM using a JEM-1400 electron microscope (Jeol Ltd., Tokyo, Japan) operating at 80 kV. The images were acquired with a Gatan Orius SC 1000B CCD camera (Gatan Inc., USA) with a resolution of 4008 × 2672 pixel image size and no binning.

### Single particle interferometric reflectance imaging

FF of individual patients, including SEV and LEV samples isolated from these FF samples, were analyzed with single particle interferometric imaging sensing (SP-IRIS) using the ExoView™ Tetraspanin kit and an ExoView™ R100 scanner (NanoView Biosciences, USA) according to the manufacturer’s instructions. Samples were diluted using the incubation buffer provided in the kit in an optimized manner based on NTA measurements (ZetaView PMX-120, Particle Metrix). FF samples were diluted to a particle concentration of 10^11^/ml, SEVs to 10^8^/ml, and LEVs to 5 × 10^9^/ml (all determined by NTA). The samples were added directly to ExoView R100 chips coated with antibodies against CD9, CD63 and CD81, respectively, and incubated at RT for 16 h. The chips were then stained using fluorescently labeled antibodies (against CD9, CD63, CD81, provided in the kit), washed, dried, and scanned. The data obtained was analyzed using the NanoViewer analysis software (NanoView Biosciences) version 3.0 with sizing thresholds set to 50 to 200 nm diameter.

The strategy for assessing the randomness of tetraspanin colocalization was based on a study by Breitwieser et al., 2022, and is illustrated in Supplementary Figure S1. In brief, the probabilities for individual events were calculated using experimental colocalization data. Individual event probabilities were then used to compute theoretical tetraspanin colocalization ratios, which were subsequently compared with experimental event ratios using a chi-square test.

### Cell culture

Human granulosa tumor-derived cell line KGN was used as a model for cell line experiments (Nishi et al., 2001). Cells were cultured in DMEM/F12 medium containing 4.5 g/l glucose, L-glutamine, and sodium pyruvate (10-013-CV, Corning) supplemented with 10% fetal bovine serum (35-089-CV, Corning) and 1% penicillin-streptomycin (Gibco, Massachusetts, USA). Cells were cultured at 37 °C in a humidified incubator with 5% CO_2_. Cells were routinely passaged after a brief exposure to 0.05% trypsin and 0.53 mM EDTA (25-051-CI, Corning).

For steroidogenesis experiments, 50,000 KGN cells were seeded on a 24-well plate in a phenol red free growth medium containing DMEM (DMEM-HXRXA, Capricorn Scientific, Hesse, Germany), 4 mM L-glutamine (GLN-B, Capricorn Scientific), 1 mM sodium pyruvate (NPY-B, Capricorn Scientific), 10% exosome-depleted fetal bovine serum (FBS, Thermo Fisher Scientific), 1% penicillin-streptomycin (Thermo Fisher Scientific), 10 μM androstenedione (LGC Standards, London, UK), and 20 ng/ml follicle stimulating hormone (Gonal-F, Merck Europe B.V., Amsterdam, Netherlands). Cells were allowed to adhere to the cell culture plate for 24 h, and were subsequently incubated with either 10^9^/ml, 10^8^/ml or 10^7^/ml SEVs, 10^8^/ml, 10^7^/ml or 10^6^/ml LEVs, or 1X DPBS (matching the highest EV sample volume) for 48h. 500 μl of cell culture medium was collected from each well, centrifuged for 5 min at 1000 g to remove cell debris. The supernatant was transferred to a new tube and stored at -80 °C until further experiments.

For transcriptomics experiments, 50,000 KGN cells were seeded on a 24-well plate in a growth medium containing DMEM/F12 (10-013-CV, Corning), 10% exosome-depleted FBS (Thermo Fisher Scientific) and 1% penicillin-streptomycin (Thermo Fisher Scientific). The cells were allowed to adhere to the cell culture plate for 48 h, then incubated with 10^9^/ml SEVs, 10^8^/ml LEVs or 1X DPBS (equal to the largest EV sample volume) in 500 µl of culture medium for 24 h. Cell culture medium was removed, cells were washed with 1X DPBS, and incubated with 700 μl of QIAzol Lysis Reagent (QIAGEN, Hilden, Germany) for 5 min. The lysates were then transferred to tubes, vortexed for 1 min, and stored at -80°C until RNA extraction.

### Cytotoxicity assay

Cytotoxicity analysis was performed using the CytoTox-Glo Cytotoxicity Assay (Promega Corporation, Wisconsin, USA) according to the manufacturer’s instructions. This assay can quantify the extracellular activity of an intracellular protease, allowing the calculation of cell viability by measuring luminescence before and after cell lysis.

10,000 KGN cells were plated into each well of a white clear-bottom 96-well plate (SPL Life Sciences, South Korea). Twenty-four hours after plating, cells were treated with either 10^6^/ml, 10^7^/ml or 10^8^/ml of LEVs, 10^7^/ml, 10^8^/ml or 10^9^/ml of SEVs or with 1X DPBS (equal to the largest EV sample volume) for 48 h. All samples were measured in triplicates with a TECAN GENios Pro plate reader (TECAN Group Ltd., Männedorf, Switzerland). Luminescence from intact cell culture and luminescence from all lysed cells were measured. The mean values of three independent experiments were used in data analysis.

### Enzyme-linked immunosorbent assay (ELISA)

Steroid hormone levels were measured from cell culture media with commercial estradiol (DRG International, New Jersey, USA), testosterone (DRG International), and progesterone (Abcam, Cambridge, UK) ELISA kits according to the manufacturer’s instructions. The limit of detection (LOD) values for the kits were 10.6 pg/ml, 0.083 ng/ml, and 0.05 ng/ml, respectively. Samples were diluted five times for estradiol and progesterone assays, and ten times for testosterone assays. The Multiskan™ FC Microplate Photometer (Thermo Fisher Scientific) was used to quantify the absorbance at a wavelength of 450 nm. The mean absorbance values for two technical replicates were calculated. A four-parameter logistic regression model was fitted to the values of measured standards, which was then used to calculate the sample concentrations. The mean values of six biological replicates were used in data analysis.

### RNA extraction

RNA extraction from isolated EVs was carried out using the miRNeasy Micro kit (QIAGEN, Hilden, Germany) following the user manual, with the modification of adding 5 µg of glycogen (R0561, Thermo Fisher Scientific) at the same time with chloroform.

RNA isolation from cell lysates included the separation of a small RNA-enriched fraction from long RNAs (>200 nt), using miRNeasy Micro kit and the RNeasy MinElute Cleanup Kit (QIAGEN) by strictly following the guidelines provided in the user manual.

RNA concentrations were measured using a Qubit 4 Fluorometer (Thermo Fisher Scientific) with either the Qubit RNA High Sensitivity Assay or the Qubit microRNA Assay (Thermo Fisher Scientific). The quality of the long RNA fraction was evaluated on an Agilent 2100 Bioanalyzer (Agilent Technologies, California, USA) with the RNA 6000 Pico kit (Agilent Technologies).

### Polyadenylated RNA sequencing and data analysis

RNA samples with RNA integrity number (RIN) values >9.0 were used for RNA sequencing. Polyadenylated RNA was isolated from 500 ng of the long RNA fraction using the NEXTFLEX Poly(A) Beads 2.0 kit (Revvity, Massachusetts, USA), which was subsequently used to prepare strand-specific RNA libraries with the NEXTFLEX® Rapid Directional RNA-Seq Kit 2.0 (Revvity) according to the manufacturer’s instructions. Library size was estimated with Agilent DNA High Sensitivity chips on the Agilent 2100 Bioanalyzer instrument (Agilent Technologies), and library concentrations were determined with the Qubit 4.0 instrument using the Qubit High Sensitivity DNA Assay (Thermo Fisher Scientific). The libraries were pooled in equimolar amounts and 150 bp paired-end sequencing was performed on the NovaSeq X platform (Illumina, California, USA). On average, 38.9 million raw reads were produced per sample (Supplementary Table 1).

Raw FASTQ reads were first assessed for their quality using FastQC version 0.11.8 (Andrews, 2010). The reads were then passed through Trimmomatic version 0.39 (Bolger et al., 2014) for quality trimming and adapter sequence removal with the parameters *LEADING:3 SLIDINGWINDOW:3:20 MINLEN:36* . The remaining read pairs were mapped to the human reference genome GRCh38 (GENCODE annotation) using STAR version 2.7.1 (Dobin et al., 2012) with default parameters. Uniquely mapped reads were counted with HTSeq-count version 0.11.2 (Anders et al., 2014), using the primary assembly annotation GTF file (version 111) obtained from Ensembl. To annotate reads that overlap with multiple features, option *intersection-nonempty* was used with HTSeq. Differential gene expression (DE) analysis was performed with DESeq2 version 1.40.2 (Love et al., 2014) in R version 4.4.1 (R Core Team, 2018). Genes with low counts were excluded from the analysis: cut-off was set at ≥ 10 normalized counts in ≥ 50% of samples. The statistical significance cut-off for DE genes was set at Benjamini-Hochberg (BH) false discovery rate (FDR) < 0.05 with no cut-off for the log_2_(fold change). PCA plot was generated with the plotPCA function in the DESeq2 package, using the top 500 most variable genes. Pathway enrichment analysis was performed for DE genes using the ShinyGO version 0.8 application tool (Ge et al., 2020) together with the Reactome database (Croft et al., 2011). Pathways with FDR < 0.05 were considered statistically significant and reported. Statistically significant Reactome terms across multiple datasets were visualized as a heatmap using the R package ComplexHeatmap (Gu et al., 2016).

### Small RNA sequencing and data analysis

Small RNA sequencing libraries were prepared from 1 ng of RNA with the QIAseq miRNA Library Kit (QIAGEN), following the manufacturer’s manual. Library size was estimated with Agilent DNA High Sensitivity chips on the Agilent 2100 Bioanalyzer instrument (Agilent Technologies). Library concentrations were determined with the Qubit High Sensitivity DNA Assay (Thermo Fisher Scientific). The libraries were pooled in equimolar amounts and single-end sequencing of 72 bp length with 10 bp dual indexing was performed on the NovaSeq X platform (Illumina). On average, approximately 30.2 million raw reads were produced per sample (Supplementary Table 2).

The raw data analysis was conducted in the QIAGEN RNA-seq Analysis Portal 5.0 using the QIAseq miRNA Library Kit analysis workflow (QIAGEN). The list of unique molecular counts of miRNAs and piRNAs was used as input for differential expression (DE) analysis with DESeq2 (version 1.40.2) in R (version 4.4.1). miRNAs and piRNAs expressed at low levels were excluded by retaining those with a sum of ≥6 unique molecular identifier (UMI) reads across SEV or LEV samples. UMI counts per million (CPM) values were calculated with the cpm function of the EdgeR package (version 4.2.2) (Robinson et al., 2010). The statistical significance cut-off for DE miRNAs and piRNAs was set at FDR < 0.05 with no cut-off for the log_2_(fold change).

The list of DE miRNAs was used as an input for the miRNA Enrichment Analysis and Annotation Tool (miEAA) (Kern et al., 2020) for over-representation analysis of Reactome pathways via miRPathDB (Backes et al., 2017). Pathways targeted by ≥ 2 miRNAs with FDR < 0.05 are reported.

mRNA targets of the DE miRNAs were predicted with the miRWalk prediction tool (Sticht et al., 2018). Only targets that intersected among miRTarBase, miRDB, and TargetScan, had target sites in their 3’-UTR region, and possessed a binding score of 1 were selected for further analysis. Cytoscape version 3.10.2 (Shannon et al., 2003) was used for plotting the network of DE miRNAs and their target genes.

### Statistical analysis

R version 4.4.1 was used to generate graphs and to perform statistical analyses. Data were analyzed using either a two-tailed Student’s t-test, Mann-Whitney U test, analysis of variance (ANOVA), or chi-square test. The specific statistical methods used in this study are described in each figure legend. Differences were considered statistically significant at *p < 0.05, **p < 0.01, and ***p < 0.001.

### Sequencing data availability

The raw data for this study have been deposited in the European Nucleotide Archive (ENA) at EMBL-EBI under accession number PRJEB83132 (https://www.ebi.ac.uk/ena/browser/view/ PRJEB83132).

## RESULTS

### Isolation and characterization of human FF EV subpopulations

To investigate the functional differences between FF EV subpopulations and their potential roles in ovarian health, we first separated EVs from FF proteins using SEC. Following this purification, we employed TFF with a 200 nm cut-off to segregate EVs by size. The molecular and morphological properties of FF EVs were characterized according to the Minimal Information for Studies of Extracellular Vesicles (MISEV) guidelines, a standard established by the International Society of Extracellular Vesicles (ISEV) (Welsh et al., 2024), before conducting any functional assays.

After SEC, we performed NTA to quantify particle concentrations, while protein concentrations were determined with BCA. Nanoparticles were detected starting from fraction 15, with a notable increase in protein concentration observed in fractions beginning at fraction 32 (Supplementary Figure S2). Thus, fractions 15-31 were pooled to ensure the effective depletion of FF proteins. The pooled EV sample was subsequently subjected to TFF, resulting in the isolation of two distinct FF EV subpopulations, here referred to as SEVs and LEVs. NTA measurements indicated minimal size overlap between the two FF EV subpopulations (Supplementary Figure S3).

We found that SEVs contained approximately 25 times more nanoparticles than LEVs, with concentrations of 1.36 × 10^11^ ± 4.18 × 10^10^ particles/ml (mean ± SEM) for SEVs and 5.43 × 10^9^ ± 1.05 × 10^9^ particles/ml for LEVs (Figure 2A). NTA measurement of particle diameters confirmed significant differences between the subpopulations, with SEVs averaging 93 ± 2 nm in diameter and LEVs 299 ± 9 nm (Figure 2B). Our isolation procedure yielded high-purity EV subpopulations, with mean particle/protein ratios of 7.93 × 10^7^ ± 1.7 × 10^7^ for SEVs and 9.08 × 10^7^ ± 1.7 × 10^7^ for LEVs (Figure 2C, Supplementary Table 3).

**Figure 2.**
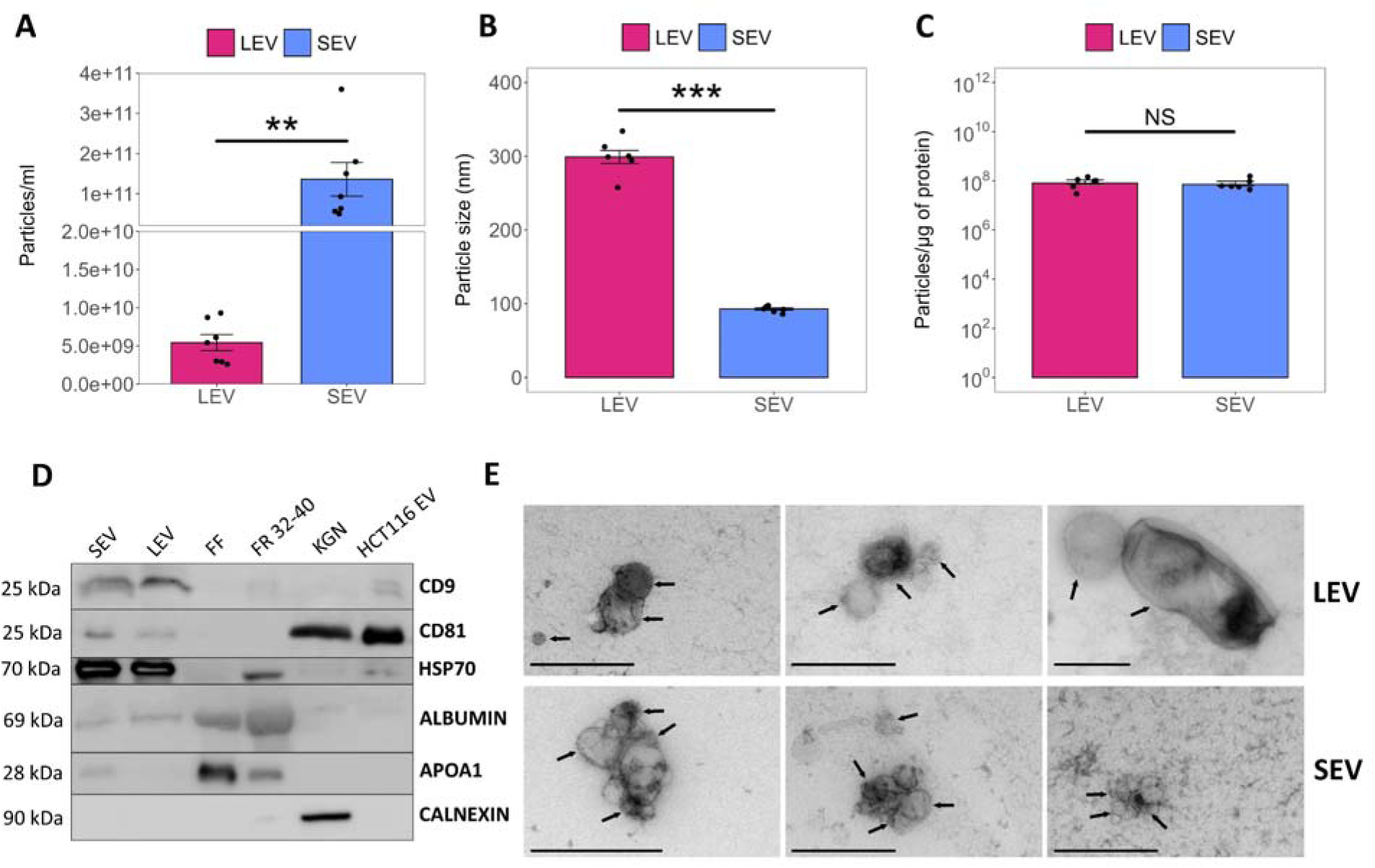
Characterization of small (SEV) and large (LEV) extracellular vesicles purified from human FF (n=6). Mean particle concentrations (**A**), diameters (**B**), and purity (**C**) of SEV and LEV samples. (**D**) Western blot analysis showing EV (CD9, CD81, and HSP70) and non-EV proteins (albumin, APOA1, and calnexin). FF and protein fractions (FR 32-40) served as a positive control for the detection of albumin and APOA1. KGN cell lysate was used as a positive control for detecting calnexin. Commercial EVs isolated from the HCT116 cell line served as a positive control for EV protein detection. (**E**) Transmission electron microscopy images of SEV and LEV samples. EVs are indicated by arrows, scale bar = 500 nm. All results are shown as mean ± SEM. Statistical differences between groups were determined by Student’s t-test and are indicated with asterisks (**p < 0.01, ***p < 0.001, NS – not significant).

To further assess EV purity, we performed Western blot analysis to detect albumin and apolipoprotein 1 (APOA1) (Figure 2D), which are abundant in FF (Kurdi et al., 2023) and often co-isolated with EVs (Simonsen, 2017). Although albumin was detected both in SEVs and LEVs, and APOA1 in SEVs, their levels were substantially reduced compared to the crude FF and the protein fractions (FR 32-40) obtained during SEC. Calnexin, an endoplasmic reticulum (ER) protein, was expectedly not detected in any EV samples but was present in the KGN cell lysate (Figure 2D).

To confirm EV enrichment in our samples, we analyzed well-established EV markers, CD9, CD81, and HSP70. Western blot revealed that CD9, CD81, and HSP70 were present in both SEV and LEV samples, while crude FF and protein fractions (FR 32-40) exhibited lower levels of these markers. Additionally, these markers were detectable in commercially produced EVs purified from the HCT116 cell line (HCT116 EV), which served as a positive control for EV marker detection (Figure 2D). TEM analysis further confirmed the presence of EV-like nanoparticles with spherical or cup-shaped morphology in both SEV and LEV samples (Figure 2E).

### Comparison of tetraspanin profiles between FF EV subpopulations

We further characterized the FF EV subpopulations using SP-IRIS to investigate the potential differences in tetraspanin colocalization between SEV, LEV and FF samples. This method captures EVs using antibodies that target specific EV proteins, followed by incubation with fluorescently labeled antibodies that bind either the same or different EV proteins. In this study, we captured and probed EVs, using antibodies against CD9, CD63, and CD81.

Our results show that both SEV and LEV samples contain all three tetraspanins – CD9, CD63, and CD81 (Figure 3). LEVs exhibit a lower percentage of CD9/CD63/CD81 colocalization compared to SEV and FF samples on the CD9-capture spot (Figure 3A), compared to FF samples on the CD63-capture spot (Figure 3B), and compared to SEV samples on the CD81-capture spot (Figure 3C). These findings suggest that SEVs are more likely to contain all three tetraspanins compared to LEVs. Conversely, LEVs were enriched in CD9-positive particles (Figure 3A) and CD9/CD81-positive particles (Figure 3C) compared to FF samples.

**Figure 3.**
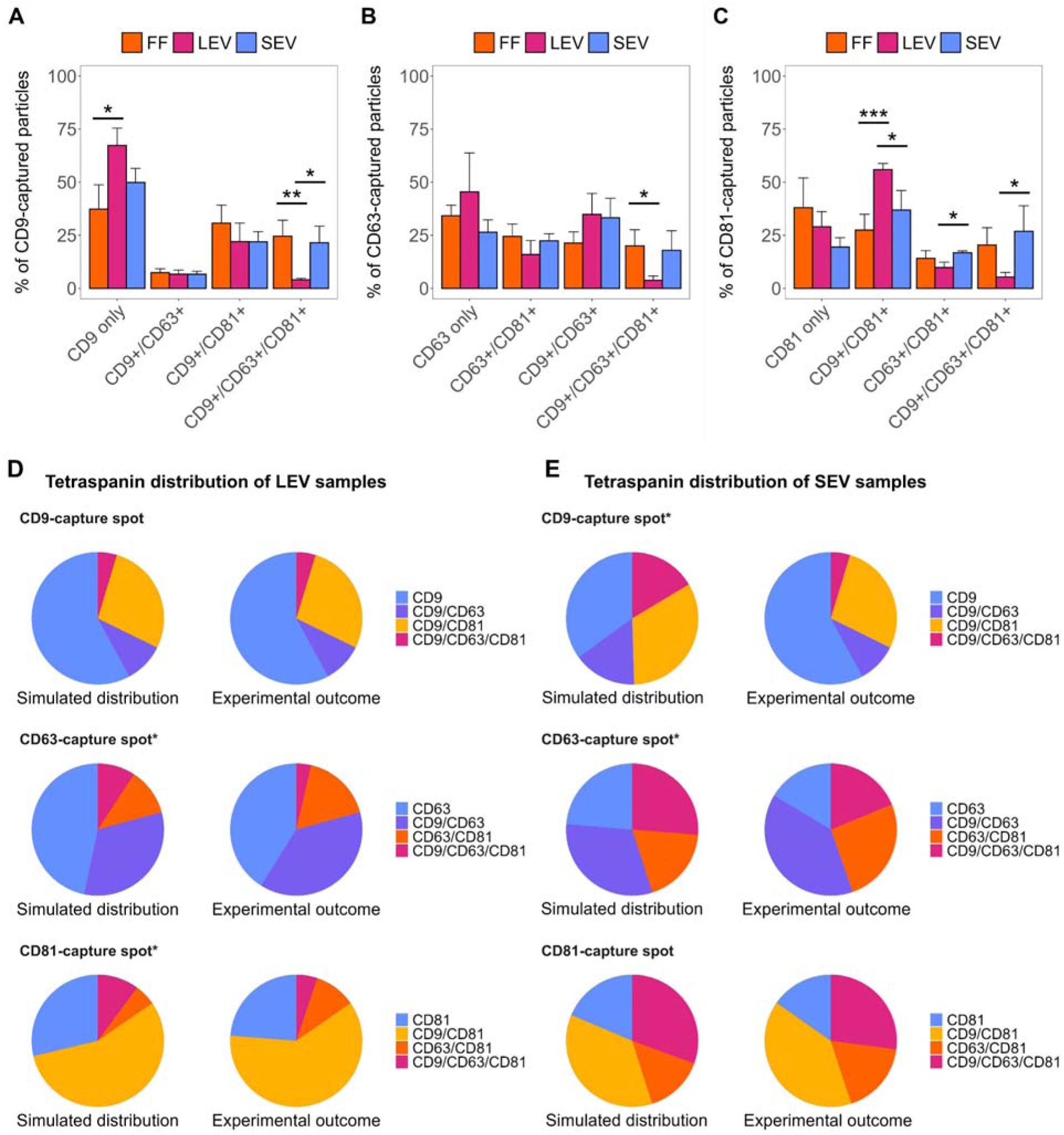
Comparison of tetraspanin profiles of SEVs, LEVs, and crude FF. SEVs (n=4), LEVs (n= 3), and FF (n=6) were loaded onto the ExoView® Tetraspanin chips and analyzed with the ExoView™ R100 scanner. Tetraspanin colocalization fractions (mean percentage of all detected EVs ± SEM) are shown for the CD9-capture spot (**A**), the CD63-capture spot, (**B**) and the CD81-capture spot (**C**). Statistical significance was determined using one-way ANOVA, followed by Tukey’s post-hoc test (*p < 0.05, **p < 0.01, ***p < 0.001). (**D**) Comparison of tetraspanin distribution between the simulated and experimental values for LEV samples and (**E**) SEV samples. Capture spots for which chi-square test between simulated and experimental colocalization ratios resulted in p < 0.05 (indicating deviation from random colocalization), are shown with asterisks (*).

Next, we assessed whether tetraspanin combinations occurred randomly or if specific tetraspanin combinations were preferentially formed in FF EVs. LEVs captured by the CD81 and CD63 antibodies displayed a statistically significant under-representation of CD9/CD63/CD81-positive particles (Figure 3D), whereas SEVs captured by the CD9 antibody showed a statistically significant over-representation of CD9/CD63/CD81 positive particles (Figure 3E) compared with simulated distributions. These results suggest that tetraspanin distribution in SEV and LEV samples does not always follow a random pattern, which may reflect differences in their biogenesis, the source of their parent cells, and/or their functions.

### The impact of FF EVs on KGN cell proliferation, viability, and steroid hormone production

In order to investigate the impact of different FF EV subpopulations on GC function, we treated the KGN cell line with varying concentrations of SEVs (10^7^, 10^8^, or 10^9^ particles/ml), LEVs (10^6^, 10^7^, or 10^8^ particles/ml), or DPBS as a control for 48 hours. The rationale for using lower concentrations of LEVs compared to SEVs in functional studies stems from their biological levels in FF being approximately a magnitude lower than those of SEVs (Supplementary Figure S3).

Cytotoxicity assays showed that neither SEVs nor LEVs significantly affected KGN cell viability (Figure 4A) or proliferation (Figure 4B) at any of the tested concentrations. Next, we assessed whether FF EV subpopulations could influence steroid hormone production in KGN cells. For this, KGN cells were treated with SEVs (10^7^, 10^8^, or 10^9^ particles/ml), LEVs (10^6^, 10^7^, or 10^8^ particles/ml), or DPBS for 24 hours, after which estradiol, progesterone, and testosterone levels were measured from the cell media by ELISA. Our results demonstrated a significant decrease in estradiol levels in KGN cells treated with the lowest SEV concentration (10^7^ particles/ml; Figure 3C), although this effect was absent at higher SEV concentrations. Additionally, treatment with the lowest LEV concentration (10^6^ particles/ml) resulted in a slight increase in progesterone production (Figure 3D). No significant changes in testosterone levels were detected in response to any of the treatments (Figure 3E).

**Figure 4.**
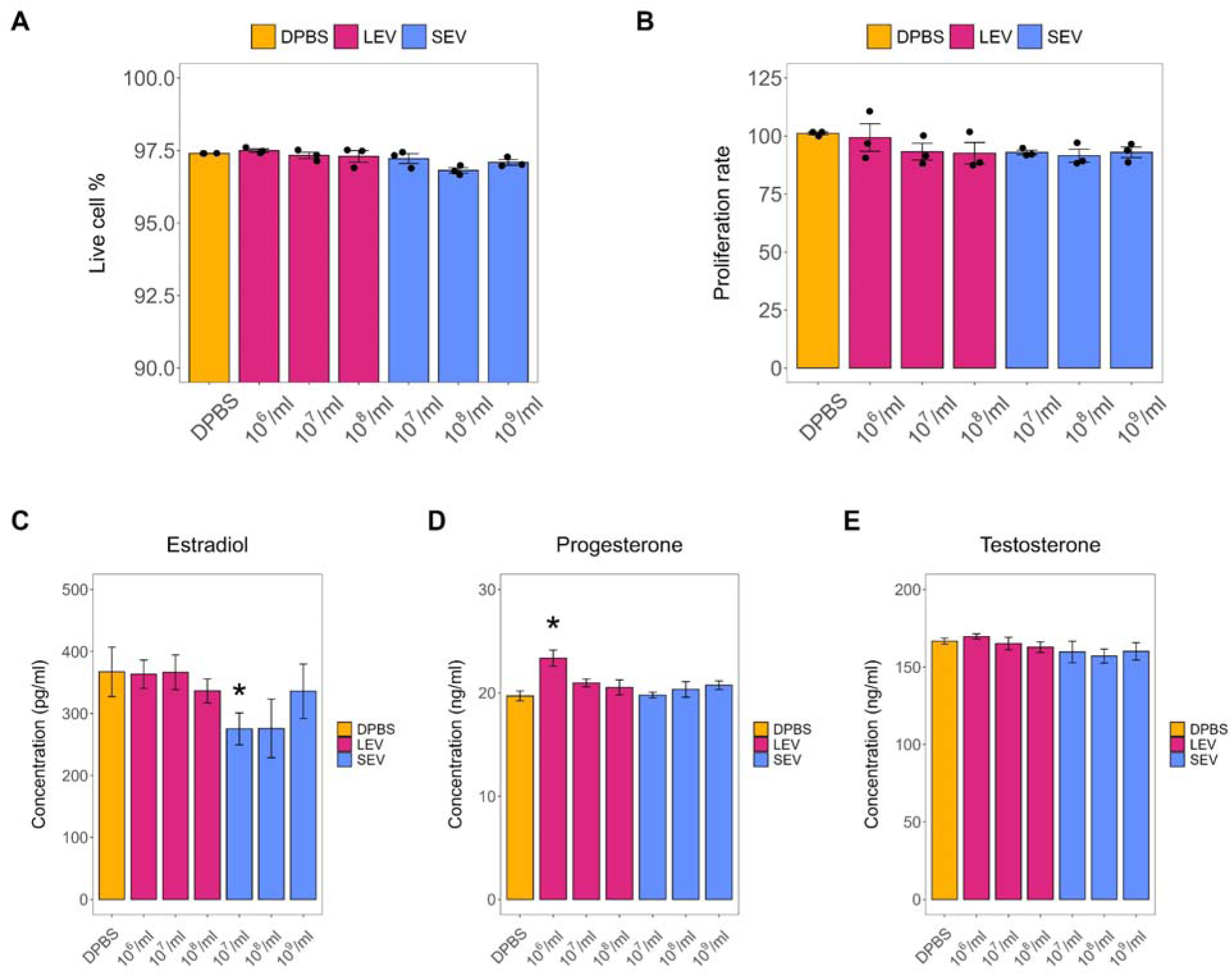
Effects of SEVs and LEVs on KGN cell viability, proliferation, and steroid hormone synthesis. KGN cells were treated with increasing concentrations of SEVs (n = 3) and LEVs (n = 3), using DPBS (n = 3) as a control, for 24 h (for steroid hormone measurements) or 48 h (for viability and proliferation measurements). After the treatment, cell viability (**A**), proliferation (**B**), and the synthesis of estradiol (**C**), progesterone (**D**), and testosterone (**E**) were assessed. All results are shown as mean ± SEM. Statistical significance was determined using the Mann-Whitney U-test (*p < 0.05).

### Differential expression of small ncRNAs in FF EV subpopulations

Aiming to better understand the role of FF EV subpopulations in ovarian intercellular communication, we analyzed their small ncRNA content through small RNA sequencing. Our analysis identified 148 miRNAs and 223 piRNAs across the analyzed samples (Supplementary Table 4).

Principal component analysis (PCA) revealed distinct expression profiles between SEV and LEV samples, with 31% of variance explained by PC1 (Figure 5A). We identified 48 differentially expressed (DE) miRNAs between SEVs and LEVs, of which 45 were upregulated in SEVs and 3 in LEVs (Figure 5B, Supplementary Table 5). Additionally, 50 DE piRNAs were identified, with 13 upregulated in SEVs and 37 in LEVs (Figure 5B, Supplementary Table 5). These results suggest that miRNAs are preferentially packaged into SEVs, while piRNAs are predominantly incorporated into LEVs by the follicular cells.

**Figure 5.**
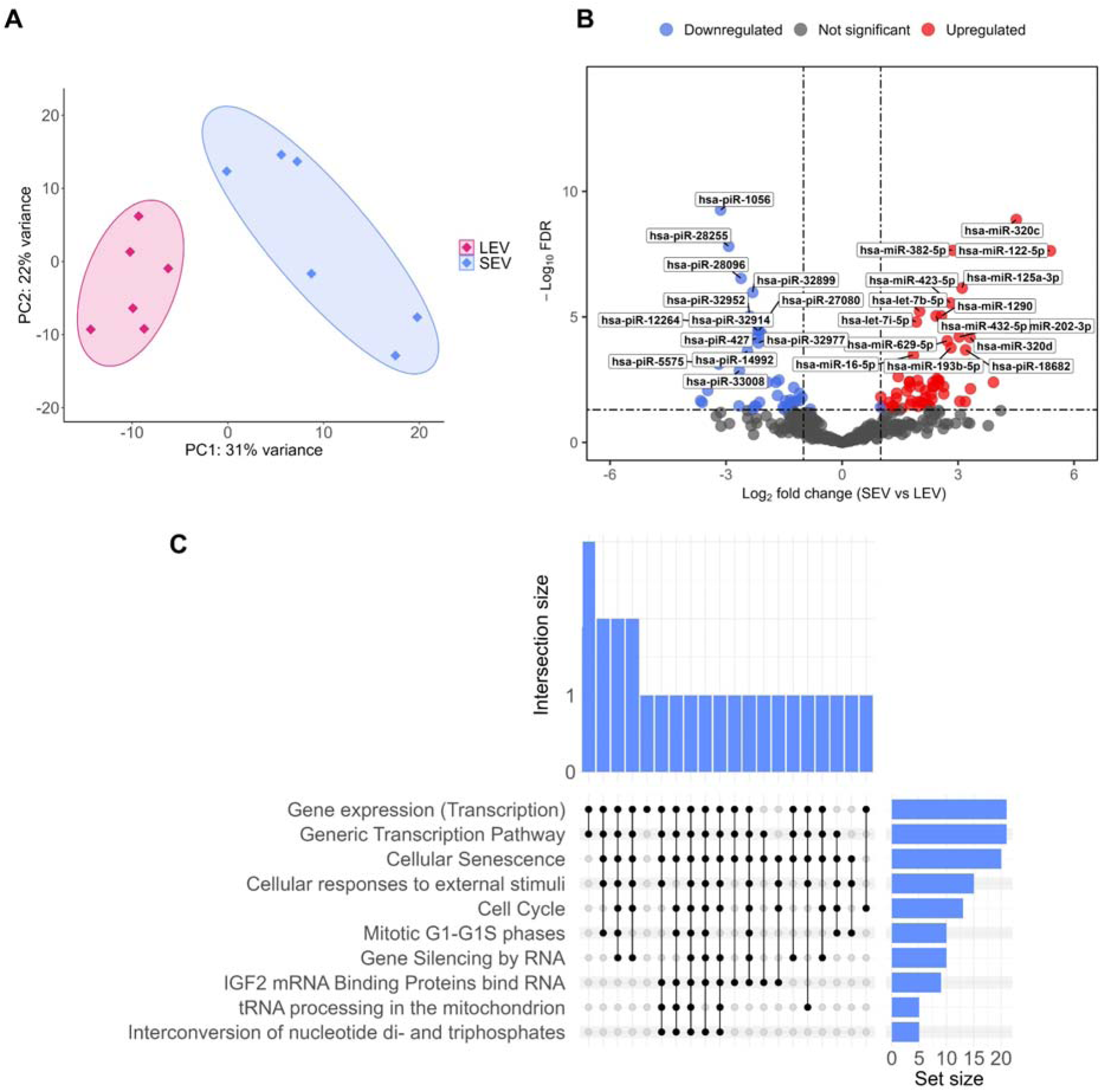
Identification of differentially expressed (DE) small noncoding RNAs between FF EV subpopulations. (**A**) PCA based on all miRNAs and piRNAs detected in SEV (n = 6) and LEV samples (n = 6). (**B**) DE miRNAs and piRNAs between SEV and LEV samples. RNAs that are more abundant (log_2_ fold change > 1) in SEVs are indicated by red dots, whereas those more abundant (log_2_ fold change < -1) in LEVs are indicated by blue dots. (**C**) Top 10 enriched pathways for DE miRNAs identified via miEAA over-representation analysis using the Reactome database. Interaction size represents the number of shared miRNAs among pathways. Set size represents the number of DE miRNAs annotated to a specific pathway.

To explore the biological processes potentially influenced by DE miRNAs, we performed over-representation analysis (ORA) with miEAA to identify enriched Reactome terms. ORA highlighted pathways related to generic transcription, cellular senescence, insulin-like growth factor-2 mRNA binding proteins (IGF2BPs), the cell cycle, cellular responses to external stimuli, and tRNA processing (Figure 5C). Additionally, pathways associated with signaling by the TGF-β receptor complex, estrogen receptor (ESR), WNT signaling, TP53 regulation of metabolic genes, Ca^2+^ signaling, and TNF signaling were identified (Supplementary Table 6). The upregulation of miRNAs associated with these pathways in SEVs suggests that small FF EVs may influence critical cellular processes in target cells.

### SEVs trigger significant changes in KGN cell gene expression

Our next objective was to determine whether FF EV subpopulations differentially influence the transcriptome of KGN cells. To address this, KGN cells were treated with the highest achievable concentration of SEVs (10^9^ particles/ml), LEVs (10^8^ particles/ml), or DPBS as a control for 24 hours, followed by RNA-sequencing. PCA of gene expression data revealed that LEV-treated cells clustered closely with DPBS-treated cells, while SEV-treated cells clustered more distinctly (Figure 6A).

**Figure 6.**
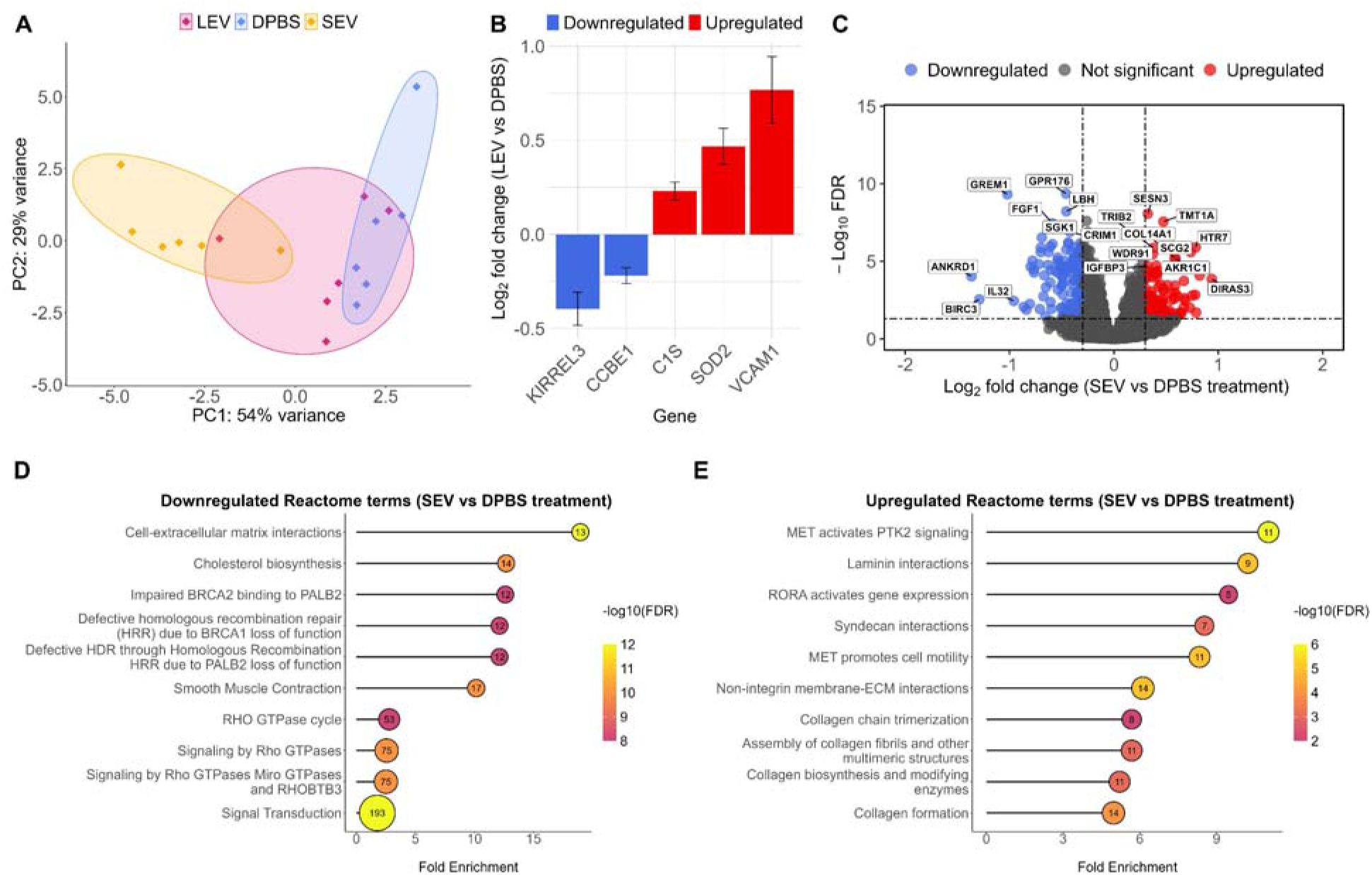
Identification of differentially expressed genes (DEGs) in KGN cells treated with SEVs (n = 6), LEVs (n = 6), or DPBS (n = 6). (**A**) PCA of mRNA expression data from KGN cells treated with SEVs, LEVs, or DPBS. (**B**) Log_2_ fold change for statistically significant (FDR < 0.05) DEGs between LEV and DPBS treatments (mean ± SEM). (**C**) Volcano plot showing DEGs upon SEV treatment, compared to DPBS-treatment. Red dots represent upregulated DEGs (log_2_ fold change > 0.3), and blue dots represent downregulated DEGs (log_2_ fold change < -0.3) with FDR < 0.05. (**D**) Top 10 Reactome terms (FDR < 0.05) enriched with downregulated genes between SEV and DPBS treatments. (**E**) Top 10 Reactome terms (FDR < 0.05) enriched with upregulated genes between SEV and DPBS treatments. The number of DEGs observed by comparing SEV vs DPBS in each pathway is noted within circles in D and E.

LEV treatment induced changes in the expression of five genes in KGN cells: KIRREL3 and CCBE1 were downregulated, while VCAM1, C1S, and SOD2 were upregulated (Figure 6B, Supplementary Table 7). In contrast, SEV treatment caused extensive transcriptomic changes, with 692 genes upregulated and 904 genes downregulated (Figure 6C, Supplementary Table 8). Enrichment analysis for Reactome pathways revealed that genes downregulated by SEV treatment were enriched in pathways related to cholesterol biosynthesis, cell-ECM interactions, homologous recombination repair, and Rho GTPase signaling (Figure 6D, Supplementary Table 9). In contrast, genes upregulated by SEV treatment were primarily associated with extracellular matrix (ECM) remodeling, including laminin and syndecan interactions, collagen biosynthesis, non-integrin membrane-ECM interactions, degradation of ECM, ECM proteoglycans, as well as MET-activated PTK2 signaling (Figure 6E, Supplementary Table 10).

To investigate SEV miRNA and KGN mRNA interactions upon incubation with SEVs, we predicted target genes of miRNAs upregulated in SEVs using miRWalk. Since miRNAs primarily mediate gene silencing (Quévillon Huberdeau & Simard, 2019), we compared the predicted miRNA targets with genes downregulated in SEV-treated cells and identified 74 miRNA-mRNA interactions (Supplementary Figure S4, Supplementary Table 11). The largest interaction network involved hsa-let-7b-5p and its target genes, which regulate cell cycle progression (CCND1, PDGFB) and cytoskeletal dynamics (FARP1, UTRN, FIGN, NAP1L1).

### SEV and LEV treatments alter small ncRNA levels in KGN cells

To further characterize the impact of FF EVs on the KGN cell transcriptome, we performed small RNA sequencing of KGN cells treated with the highest achievable concentration of SEVs (10^9^ particles/ml), LEVs (10^8^ particles/ml), or DPBS as a control for 24 hours. PCA revealed distinct clustering for all treatments (Figure 7A), highlighting the differential effects of SEVs and LEVs on small ncRNA levels in KGN cells.

**Figure 7.**
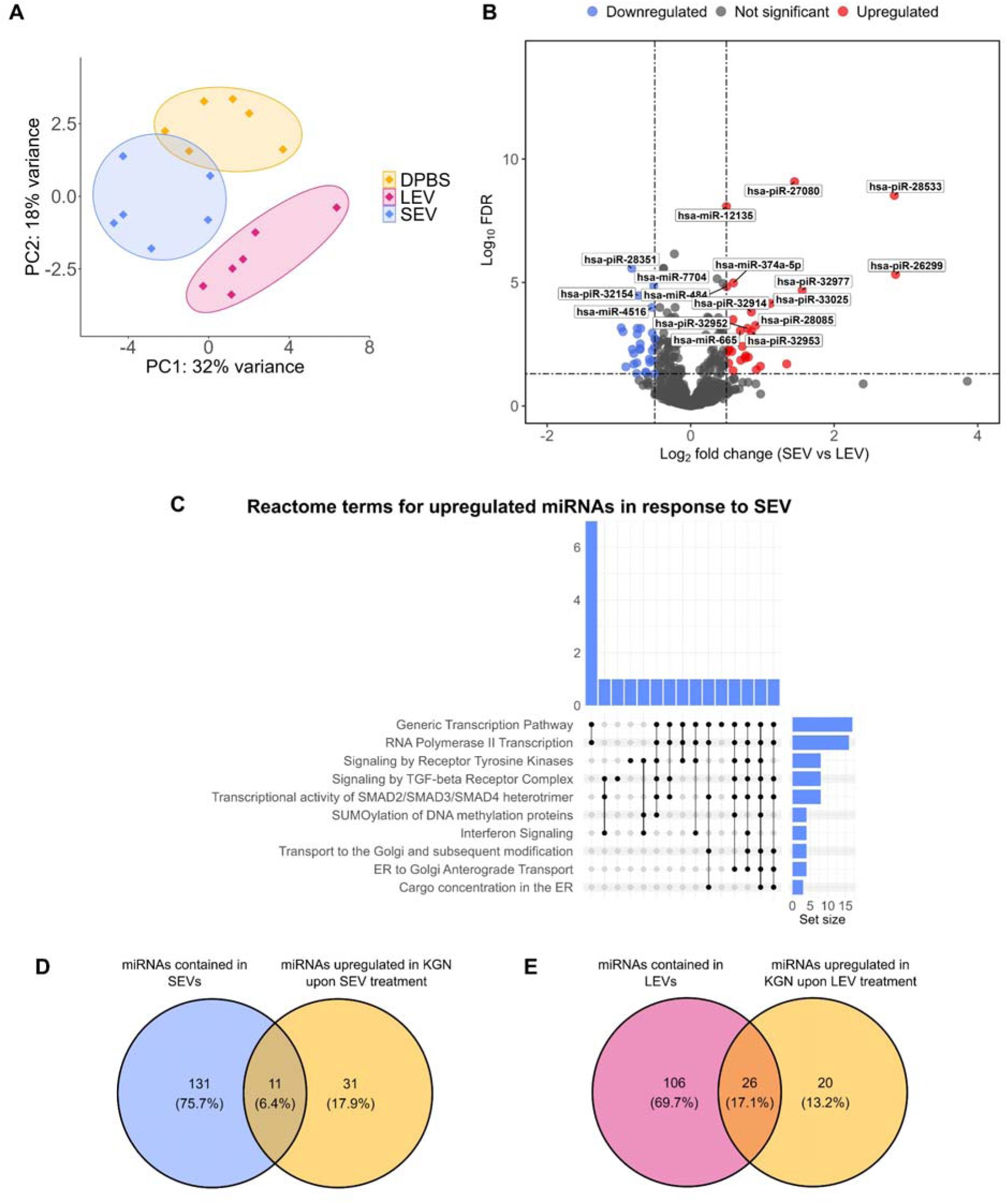
Identification of DE ncRNAs in KGN cells treated with SEVs (n = 6), LEVs (n = 6), or DPBS (n = 6). (**A**) PCA based on piRNA and miRNA expression levels of KGN cells treated with SEVs, LEVs, or DPBS. (**B**) Volcano plot showing DE ncRNAs between SEV and LEV treatments. Red dots represent upregulated ncRNAs in SEVs (log_2_ fold change > 0.3), and blue dots represent upregulated ncRNAs in LEVs (log_2_ fold change < -0.3) with FDR < 0.05. (**C**) Top 10 Reactome terms for upregulated miRNAs in response to SEV treatment. Reactome terms were identified via miEAA over-representation analysis. Interaction size represents the number of shared miRNAs. Set size represents the number of miRNAs annotated to a specific pathway. (**D**) Overlap between miRNAs upregulated in KGN cells upon SEV treatment and miRNAs contained in SEVs. (**E**) Overlap between miRNAs upregulated in KGN cells upon LEV treatment and miRNAs contained in LEVs.

DE analysis between SEV- and LEV-treated samples identified 159 DE ncRNAs, including 88 DE miRNAs and 71 piRNAs (Figure 7B, Supplementary Table 12). Of the DE miRNAs, 42 were upregulated and 46 downregulated in SEV-treated cells; while of the DE piRNAs, 27 were upregulated and 44 downregulated in SEV-treated cells (Supplementary Table 12).

ORA using the Reactome database revealed distinct functional associations for miRNAs upregulated in SEV- and LEV-treated cells. miRNAs upregulated upon LEV treatment were enriched in pathways related to IGF2BPs (FDR = 6.09 × 10^−3^) and PTEN regulation (FDR = 0.0176), whereas miRNAs upregulated upon SEV treatment were associated with pathways related to generic transcription, receptor tyrosine kinase signaling, ER to Golgi transport, TGF-β signaling, and Smad2/Smad3/Smad4 transcriptional activity (Figure 7C, Supplementary Table 13).

We examined whether miRNAs present in FF EVs were upregulated in KGN cells, suggesting their potential delivery via EVs. Among the 42 miRNAs upregulated upon SEV treatment in KGN cells, 11 were present in SEVs (Figure 7D), and among the 46 miRNAs upregulated upon LEV treatment, 26 were detected in LEVs (Figure 7E). Notably, there was no overlap between miRNAs potentially delivered by SEVs and LEVs. For piRNAs, 20 of the 27 piRNAs upregulated in SEV-treated cells were present in SEVs (Supplementary Figure S5), while 28 of the 44 upregulated piRNAs in LEV-treated cells were present in LEVs (Supplementary Figure S6).

### Integration of gene ontology datasets reveals central pathways modulated by SEVs

To identify key pathways influenced by SEVs, we integrated pathway enrichment results from four independent analyses: DE miRNAs between SEVs and LEVs (45 of 48 DE miRNAs were upregulated in SEVs), downregulated mRNAs in SEV-treated GCs, upregulated mRNAs in SEV-treated GCs and upregulated miRNAs in SEV-treated GCs. This systems-level approach identified overlapping pathways, highlighting their robustness and biological significance.

The analysis identified transcriptional regulation as a central target of SEVs, with both the generic transcription pathway and RNA polymerase transcription consistently enriched across all four datasets (Figure 8). Notably, SEV treatment led to the upregulation of POLR2A (Supplementary Table 8), the largest subunit of RNA polymerase II, and GTF2F1, a component of the RNA polymerase II preinitiation complex, suggesting that SEVs may enhance transcriptional efficiency or activity in GCs.

**Figure 8.**
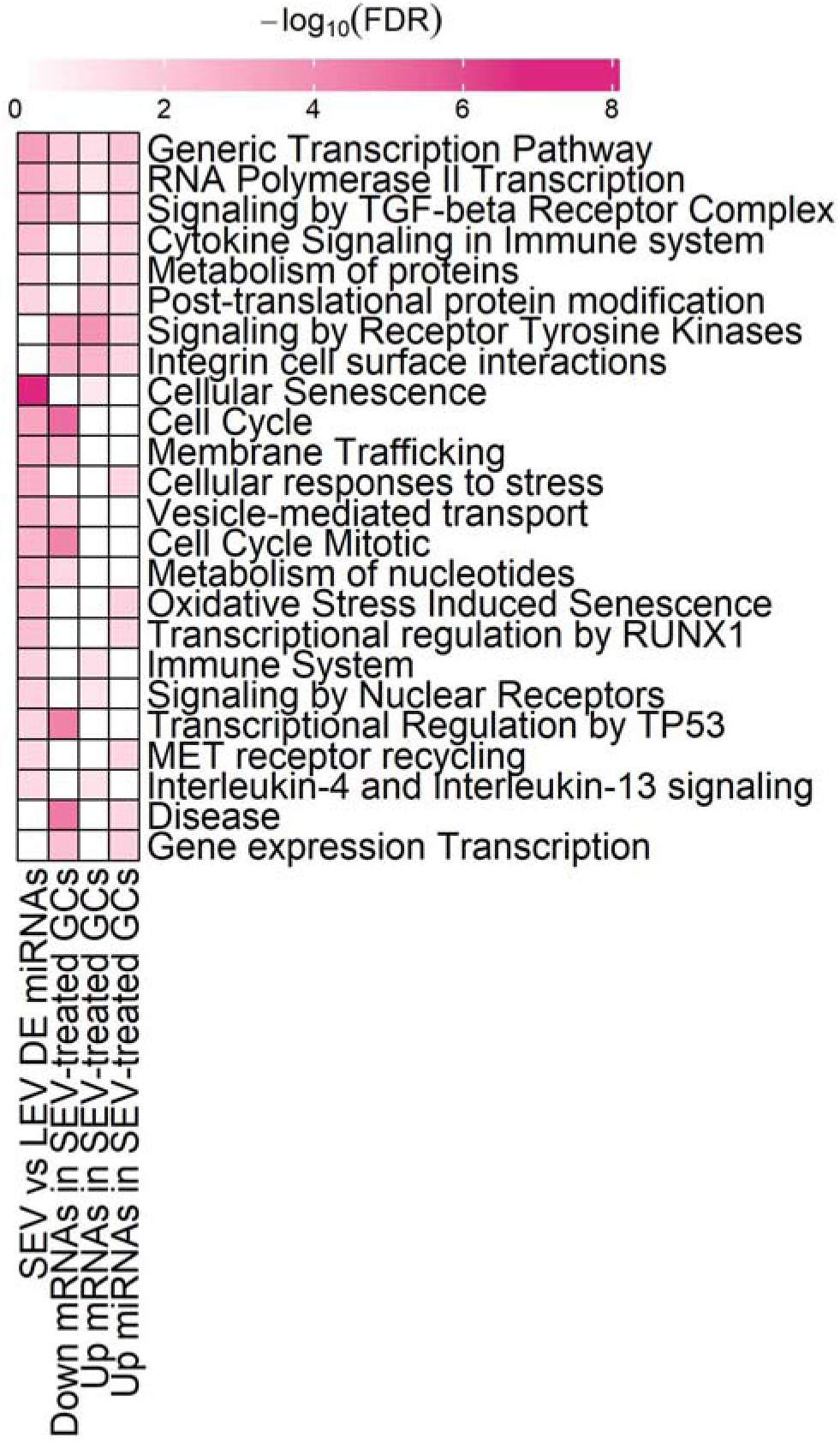
Heatmap of statistically enriched Reactome terms associated with SEVs: DE miRNAs predominantly upregulated in SEVs compared to LEVs, downregulated mRNAs in SEV-treated GCs, upregulated mRNAs in SEV-treated GCs and upregulated miRNAs in SEV-treated GCs. Color intensities represent the -log_10_(FDR) values for each Reactome term.

In addition to transcriptional regulation, six pathways were consistently enriched across three datasets, underscoring their importance in SEV-mediated processes. Among these, signaling by the TGF-β receptor complex, which is known for its roles in follicular development, GC proliferation and oocyte maturation (Knight & Glister, 2006), emerged as a key pathway. The enrichment of the cytokine signaling pathway reflects the role of SEVs in immune regulation within the follicle, while pathways related to protein metabolism and post-translational protein modifications emphasize the potential role of SEVs in fine-tuning protein activity, turnover, and stability in GCs. The consistent enrichment of receptor tyrosine kinase signaling points to the regulation of cell survival and differentiation, and integrin cell surface interactions link SEVs to ECM remodeling and cell-ECM communication.

Pathways enriched across two datasets further reinforce the multifunctional roles of SEVs, particularly in cellular stress responses and cell cycle regulation. Cellular stress pathways, including oxidative stress-induced senescence and cellular responses to stress, indicate SEV-driven mechanisms that may help protect GCs from cellular damage and maintain their viability. Taken together, these findings offer a systems-level understanding of SEV-mediated intercellular communication and suggest that SEVs play a multifaceted role in reprogramming GC functions, particularly through the modulation of transcriptional activity and cellular signaling.

## DISCUSSION

FF-derived EVs play a crucial role in facilitating intercellular communication between oocytes and somatic cells of the ovarian follicle. To better understand their distinct contributions to the follicular microenvironment, small and large EVs were studied separately, as they may differ in molecular composition and biological activities. By isolating and characterizing these EV subpopulations, we aimed to gain a clearer understanding of the GC function, including viability, proliferation, steroidogenesis, and gene expression.

Our results demonstrated that SEVs are significantly more abundant than LEVs in FF, consistent with earlier reports by Neyroud et al., 2022 and Franz et al., 2016, who similarly observed higher abundance of SEVs among FF EVs. Single-EV tetraspanin profiling revealed a non-random distribution of tetraspanins, with LEVs containing a lower proportion of CD9/CD63/CD81-positive particles compared to SEVs and crude FF. These differences likely reflect distinct biogenesis pathways and functional roles of SEVs and LEVs, emphasizing the need for further investigation into how these differences influence EV uptake and intracellular trafficking.

While previous studies in bovine and human models reported that FF EVs promote GC proliferation (Couty et al., 2024; Hung et al., 2017; Ying et al., 2023), our findings showed no such effect in SEV- or LEV-treated KGN cells. This discrepancy may arise from differences in EV isolation methods, biological variations between species, and/or experimental conditions. Importantly, we employed size-exclusion chromatography, a method known to reduce FF protein contamination compared to ultracentrifugation (UC) (Soares et al., 2023), which likely improved the purity of our EV preparations. This raises the possibility that proliferative effects reported in UC-based studies were partly influenced by residual FF proteins rather than EV internal cargo. Interestingly, while both our results and those of Couty et al., 2024 showed no effect under normal physiological conditions, Zhou et al., 2023 demonstrated that FF EVs reduced GC apoptosis under stress conditions in a PCOS-like model. This suggests that FF EVs may serve protective functions under stress conditions, while exhibiting minimal effects under normal physiological conditions.

Consistent with animal studies showing that FF EVs modulate steroidogenesis (Ying et al., 2023; Yuan et al., 2021), we observed dose-specific effects of SEVs and LEVs on steroid hormone production. SEVs reduced estradiol levels, whereas LEVs increased progesterone synthesis at the lowest tested EV concentrations, aligning with the physiological steroidogenic shift following the LH surge. This transition, marked by declining estradiol production and increased progesterone synthesis (Maman et al., 2023), is crucial for GC luteinization, suggesting that SEVs and LEVs isolated from preovulatory follicles may play complementary roles in regulating this process. At the highest EV doses, steroidogenesis-related gene expression was largely unaffected, apart from the 2.3-fold upregulation of *CYP19A1* in SEV-treated cells. These findings raise the possibility that changes in steroidogenesis-related gene expression occurred earlier than the 24-hour time point analyzed in this study. Further investigation is warranted to clarify how EV dose and timing influence steroidogenic pathways in GCs.

Small RNA sequencing revealed distinct ncRNA profiles in SEVs and LEVs, further highlighting their functional divergence. SEVs, enriched in miRNAs, were associated with pathways critical for GC proliferation and differentiation, including cell cycle regulation, IGF2BPs, TGF-β, WNT, TNF, TP53, and Ca^2+^ signaling. Several of these pathways have previously been identified as targets of FF EV miRNAs in both animal (da Silveira et al., 2012; Liu et al., 2024) and human (Martinez et al., 2018; Santonocito et al., 2014) studies, underscoring the conserved regulatory roles of FF EV miRNAs across species. In contrast, LEVs, enriched in piRNAs, appear to play a role in preserving genome stability, as piRNAs are essential for silencing transposable elements and maintaining genomic integrity in both germ and somatic cells (Wu et al., 2020).

These differences in ncRNA profiles were reflected in the distinct transcriptomic responses of KGN cells treated with SEVs or LEVs. LEV treatment resulted in minimal transcriptomic changes, with only five differentially expressed genes, primarily linked to immune regulation (VCAM, C1A) and oxidative stress defense (SOD2), suggesting that LEVs may play a role in maintaining cellular homeostasis. In contrast, SEV treatment induced extensive transcriptomic changes, with 1596 genes showing differential expression. SEVs upregulated genes involved in ECM remodeling and MET-activated PTK2 signaling, both of which are essential for follicular expansion and oocyte-GC communication (McGinnis & Kinsey, 2014; Vasse et al., 2024). ECM remodeling within the ovarian follicle involves the controlled synthesis and degradation of structural components like collagens and laminins (Irving-Rodgers & Rodgers, 2005). PTK2, activated through MET signaling, regulates focal adhesions — specialized cellular structures that transmit mechanical forces and signaling cues between the ECM and interacting cells, essential for tissue remodeling. These findings underscore the role of SEVs in facilitating the structural reorganization necessary for ovulation, aligning with previous studies highlighting FF EV roles in ECM formation (de Almeida Monteiro Melo Ferraz et al., 2020) and ECM-receptor interactions (Martinez et al., 2018).

SEV treatment led to the downregulation of genes associated with Rho GTPases, cell cycle regulation, cholesterol biosynthesis, and homologous recombination repair (HRR). The suppression of Rho GTPases, which regulate cytoskeletal organization and mitosis (Chircop, 2014), likely reflects necessary cytoskeletal and cell cycle adaptations for GC differentiation during luteinization, a process characterized by cell cycle exit (Robker & Richards, 1998). Furthermore, the downregulation of cholesterol biosynthesis genes may represent a metabolic shift favoring cholesterol uptake rather than synthesis to meet the demands of hormone production. Alternatively, it may reflect underlying fertility challenges, as reduced cholesterol biosynthesis is linked to diminished ovarian reserve (Yang et al., 2022). Similarly, suppression of HRR genes, such as BRCA2 and RAD51, indicates potential genomic instability, which is a recognized contributor to infertility (Dunce & Davies, 2024). Collectively, these findings suggest SEVs influence both the structural and metabolic states of GCs, supporting their transition during luteinization.

Analysis of ncRNA profiles in FF EV-treated GCs further distinguished the effects of SEVs and LEVs. LEV-treated GCs exhibited upregulated miRNAs linked to IGF2BPs and PTEN regulation, pathways associated with GC survival. Elevated levels of PTEN, a negative regulator of the PI3K-AKT pathway, has been implicated in impaired follicular development and increased GC apoptosis (Yao et al., 2021). Although LEV supplementation itself did not alter PTEN mRNA levels after 24 h of incubation, miRNAs targeting PTEN could repress its translation, suggesting that LEVs may contribute to GC survival and cellular homeostasis. Conversely, SEV-treated GCs exhibited upregulated miRNAs associated with transcription, TGF-β signaling, and Smad transcriptional activity. These pathways align with the functional roles of SEV-enriched miRNAs, suggesting SEVs may impact GCs through direct miRNA delivery, but also through secondary transcriptional changes.

The integration of pathway enrichment analyses further strengthens these findings by revealing recurring pathways influenced by SEVs. Modulation of transcriptional pathways by SEVs is particularly noteworthy, as GCs must undergo extensive transcriptional changes to adapt to the dynamic physiological demands of follicular development and luteinization. Another key pathway impacted by SEVs is TGF-β signaling, which was significantly downregulated in SEV-treated GCs. TGF-β is known to suppress oocyte maturation (Coskun & Lin, 1994) and maintain meiotic arrest (Yang et al., 2019) during early follicular development, and its suppression after the LH surge is essential for meiotic resumption. SEV-mediated downregulation of TGF-β signaling may also support GC luteinization by alleviating its inhibitory effects on *STAR* expression and progesterone synthesis (Fang et al., 2014; Zheng et al., 2009). Furthermore, TGF-β regulates ECM remodeling and GC apoptosis during the periovulatory period (Wang et al., 2019). By regulating this pathway, SEVs may facilitate ECM reorganization and inhibit apoptosis to ensure GC survival during luteinization.

Similarly, the enrichment of the integrin cell surface interactions pathway highlights the role of SEVs in ECM remodeling. Integrins, which mediate cell-ECM adhesion, are critical for cellular communication and structural organization (Anderson et al., 2013). Among the upregulated genes in this pathway, fibronectin (FN1) stands out as a key ECM glycoprotein that binds integrins and supports GC adhesion. The SEV-induced upregulation of fibronectin is particularly significant, as it has been shown to promote GC luteinization and cumulus expansion (Kitasaka et al., 2018). The concurrent downregulation of genes encoding collagens and specific integrin subunits suggests a regulated loosening of cell-ECM adhesion, enabling the dynamic remodeling required during follicular maturation.

Finally, cytokine signaling emerged as another pathway influenced by SEVs. Cytokines, particularly interleukins (ILs), are well-known for their roles in immune regulation and inflammation. However, the ovarian follicle itself is a site of inflammatory-like processes, where ovarian cells both produce and respond to cytokines. Among these, IL-1 has been shown to regulate ovulation-associated events, including the production and activation of proteolytic enzymes, such as prostaglandins, which mediate follicle rupture (Gérard et al., 2004; Peterson et al., 1993). The upregulation of IL1R1, the receptor for IL-1, in SEV-treated GCs suggests that SEVs may enhance IL-1 signaling necessary for follicle wall breakdown and oocyte release.

Despite the significant findings, this study has several limitations. First, the inclusion of patients undergoing ovarian stimulation may have influenced the FF EV profile. Ovarian stimulation is known to alter FF miRNA profiles (Noferesti et al., 2015), potentially introducing variability into our findings. Future studies utilizing patient material collected during natural cycles would provide a clearer understanding of how ovarian stimulation affects EV composition. However, such samples are not available from the majority of infertility clinics. Secondly, while the KGN cell line is advantageous for reducing variability across experiments, it is a tumor-derived GC-like model that may not fully replicate the behavior of healthy GCs *in vivo*. Further research involving primary human GCs would provide more physiologically relevant insights into FF EV-mediated mechanisms. Finally, the SEV and LEV populations were not exclusively separated based on the TEM visualization. However, a complete separation would be very difficult to achieve.

Nevertheless, this study has notable strengths. By employing rigorous EV isolation and characterization protocols, including single-EV profiling and small RNA sequencing, we ensured high purity and size-specific enrichment of SEVs and LEVs. Additionally, the integration of transcriptomic and ncRNA analyses enabled us to uncover nuanced functional and molecular distinctions between FF EV subpopulations, addressing gaps in prior research that often examined pooled FF EVs.

Taken together, this study highlights the functional and molecular differences between SEVs and LEVs in human FF, emphasizing their distinct contributions to ovarian function. SEVs, more abundant and enriched in miRNAs, were found to regulate pathways related to transcription, TGF-β signaling, ECM remodeling, and cell cycle, suggesting their role in driving structural and functional changes during follicular development. LEVs, less abundant and enriched in piRNAs, were linked to IGF2BPs and PTEN regulation, emphasizing their role in maintaining genomic stability and cellular homeostasis. These findings advance our understanding of intercellular communication within the ovarian follicle and underscore the importance of considering EV heterogeneity in ovarian biology studies.

## Supporting information

Graphical abstract

## GEOLOCATION INFORMATION

59.39739419137438, 24.662140692464533

## ACKNOWLEDGEMENTS

We are thankful to the staff of Nova Vita Clinic and East Tallinn Central Hospital for recruiting study participants and to the participants themselves for the donation of their samples for this study.

The authors acknowledge support from the National Genomics Infrastructure in Stockholm funded by Science for Life Laboratory, the Knut and Alice Wallenberg Foundation and the Swedish Research Council. We also thank NAISS/Uppsala Multidisciplinary Center for Advanced Computational Science for assistance with massively parallel sequencing and access to the UPPMAX computational infrastructure. We acknowledge the Electron Microscopy Unit of the Institute of Biotechnology, University of Helsinki, for providing the microscopy facilities.

## AUTHOR CONTRIBUTIONS

**Inge Varik** : Formal analysis; Methodology; Investigation; Data curation; Writing – original draft, Visualization. **Katariina Johanna Saretok** : Methodology; Writing – review and editing. **Kristine Rosenberg:** Resources; Writing – review and editing. **Ileana Quintero:** Methodology; Writing – review and editing. **Maija Puhka:** Methodology; Writing – review and editing. **Nataliia Volkova:** Methodology; Writing – review and editing. **Aleksander Trošin:** Resources; Writing – review and editing. **Paolo Guazzi:** Conceptualization; Writing – review and editing; Supervision. **Agne Velthut-Meikas:** Conceptualization; Investigation; Data curation; Writing – review and editing; Supervision; Project administration; Funding acquisition. All authors have read and agreed to the published version of the manuscript.

## DECLARATION OF GENERATIVE AI AND AI-ASSISTED TECHNOLOGIES IN THE WRITING PROCESS

During the preparation of this work the authors used ChatGPT-4o in order to improve sentence structure and correct grammar. After using this tool, the authors reviewed and edited the content as needed and take full responsibility for the content of the publication.

