## Supplementary material for "Small and large extracellular vesicles from human preovulatory follicular fluid display distinct ncRNA cargo profile and differential effect on granulosa cell line KGN": Graphical abstract

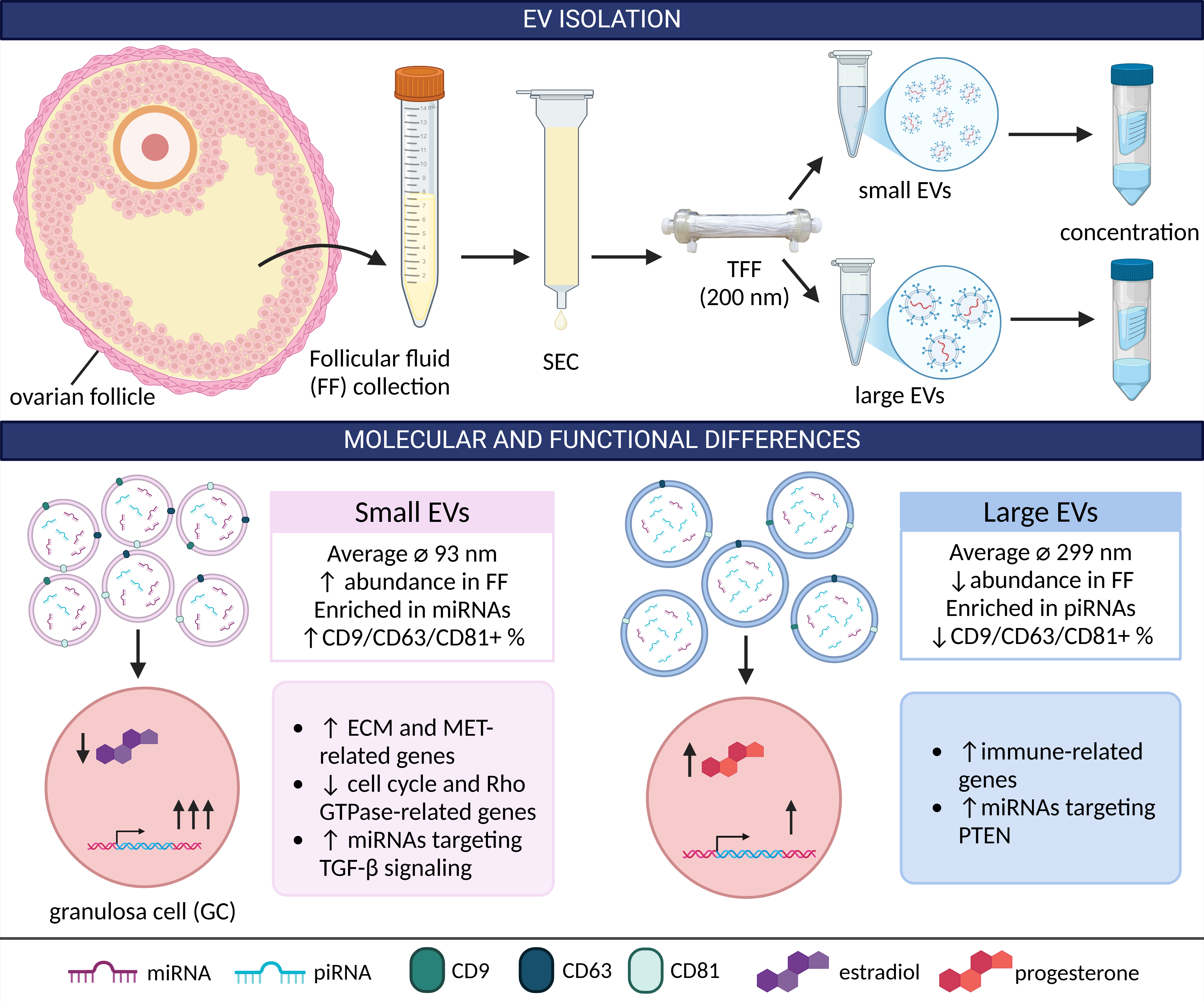

Small and large EVs isolated from human follicular fluid using SEC and TFF exhibited differences in size, abundance, tetraspanin colocalization, and short non-coding RNA cargo. These EV subpopulations had divergent effects on the KGN granulosa cell line: small EVs significantly altered the expression of genes linked to the extracellular matrix, MET signaling, cell cycle, and Rho GTPases, while large EVs caused minimal gene expression changes, primarily upregulating immune-related genes. Created in BioRender. Varik, I. (2025) <https://BioRender.com/x73d811>.
