## Supplementary Figures for "Small and large extracellular vesicles from human preovulatory follicular fluid display distinct ncRNA cargo profile and differential effect on granulosa cell line KGN"

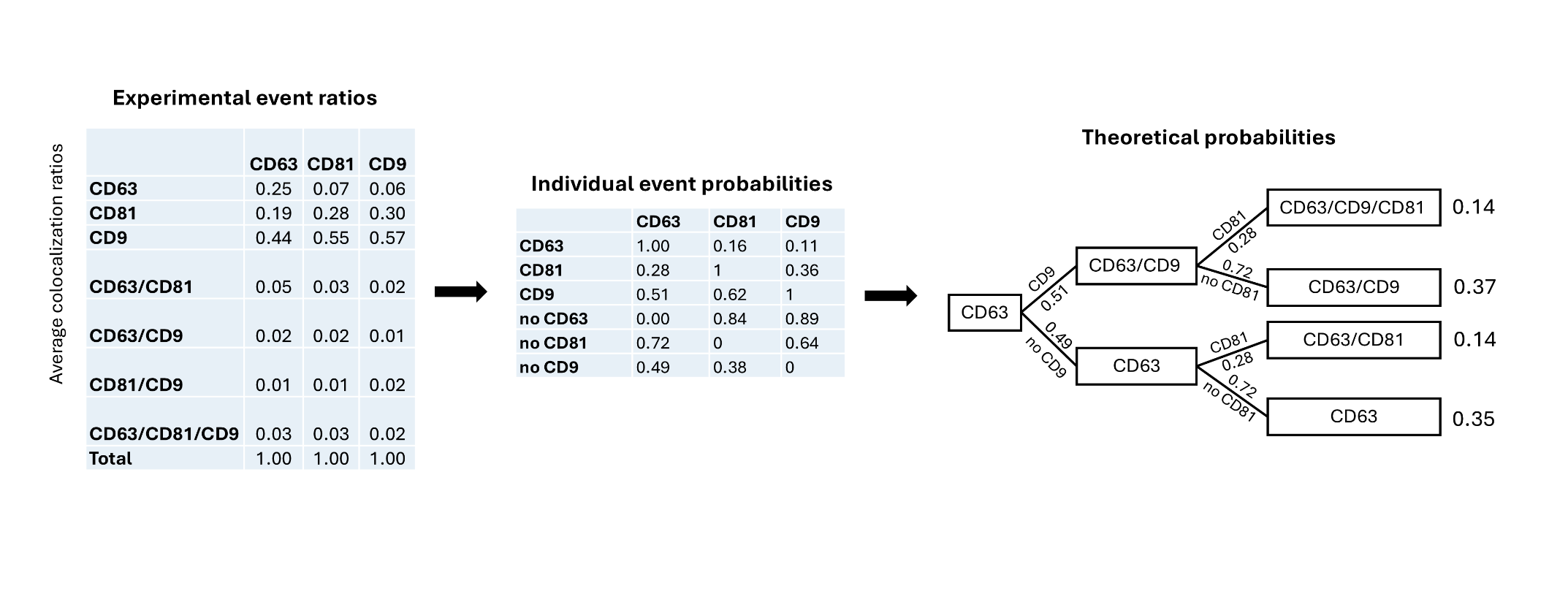

**Supplementary Figure S1.** Strategy for the calculation of tetraspanin colocalization randomness. First, the probabilities for individual events were calculated using experimental colocalization data. Individual event probabilities were then used to compute theoretical tetraspanin colocalization ratios, which were subsequently compared with experimental event ratios using a chi-square test.

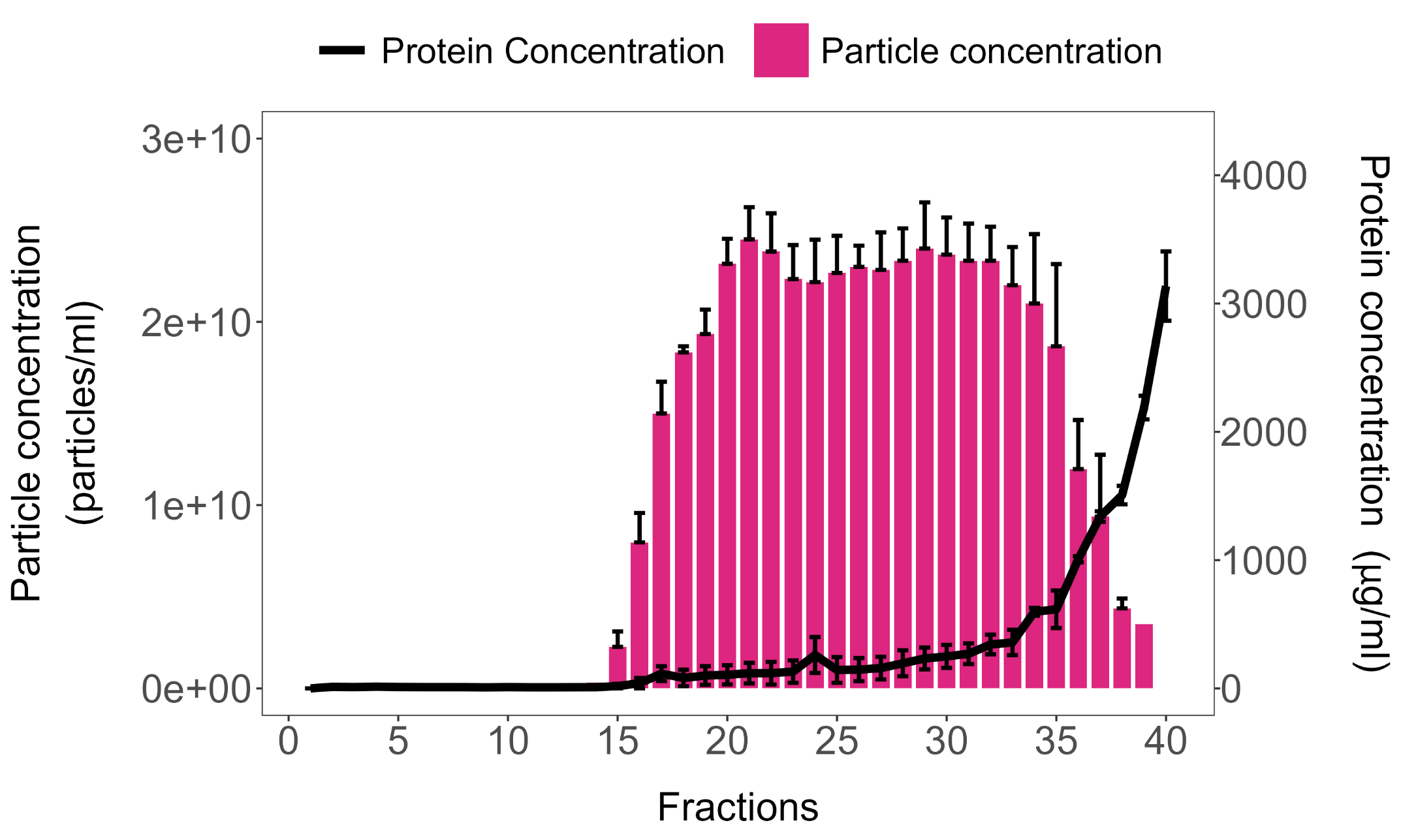

**Supplementary Figure S2.** Particle and protein concentrations for each individual fraction collected during SEC. Particle concentrations (particles/ml) were determined by NTA and protein concentrations (μg/ml) were determined by BCA (mean ± SEM, n = 3).

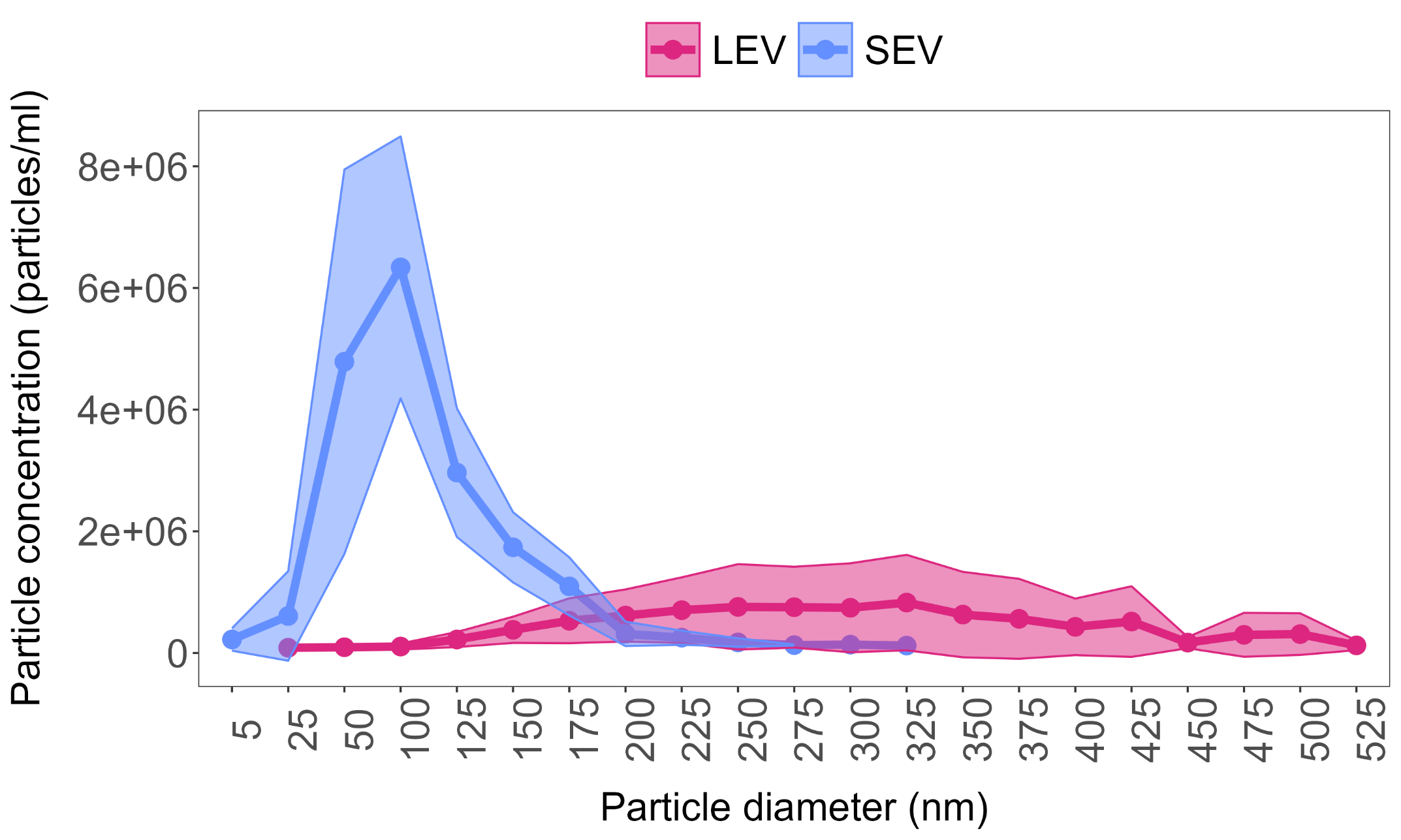

**Supplementary Figure S3.** Particle concentrations and diameters of small EVs (SEVs) and large EVs (LEVs) after isolation with TFF. Mean values are indicated by the central line, with SEM represented by the surrounding shaded area (n=3).

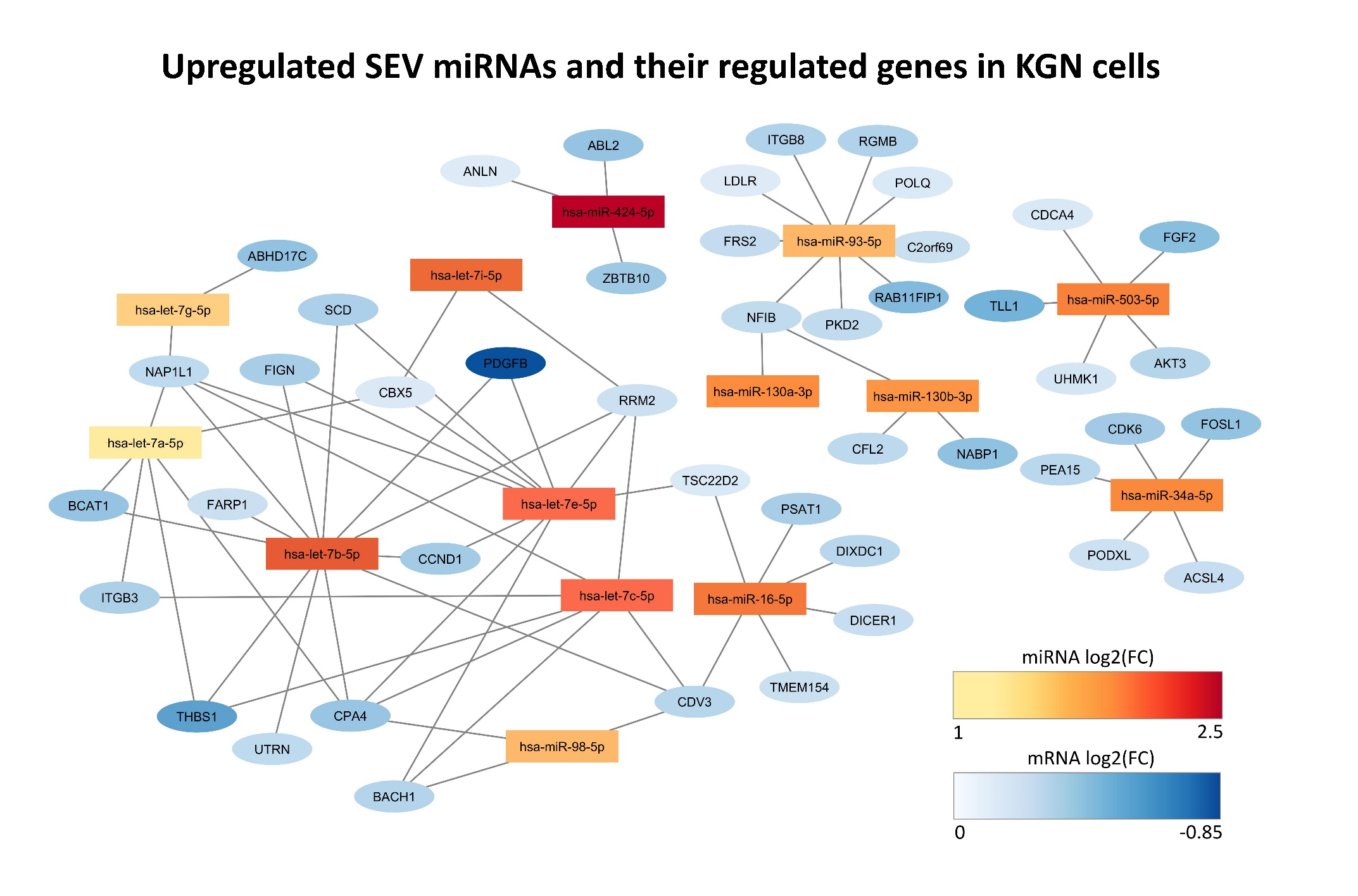

**Supplementary Figure S4.** miRNA-mRNA network illustrating upregulated miRNAs in SEVs that are predicted to downregulate several mRNAs in KGN cells. miRNAs are colored based on the fold change (FC) of differential expression (DE) between SEV and LEV samples, while mRNAs are colored based on the fold change of DE between SEV and DPBS samples.

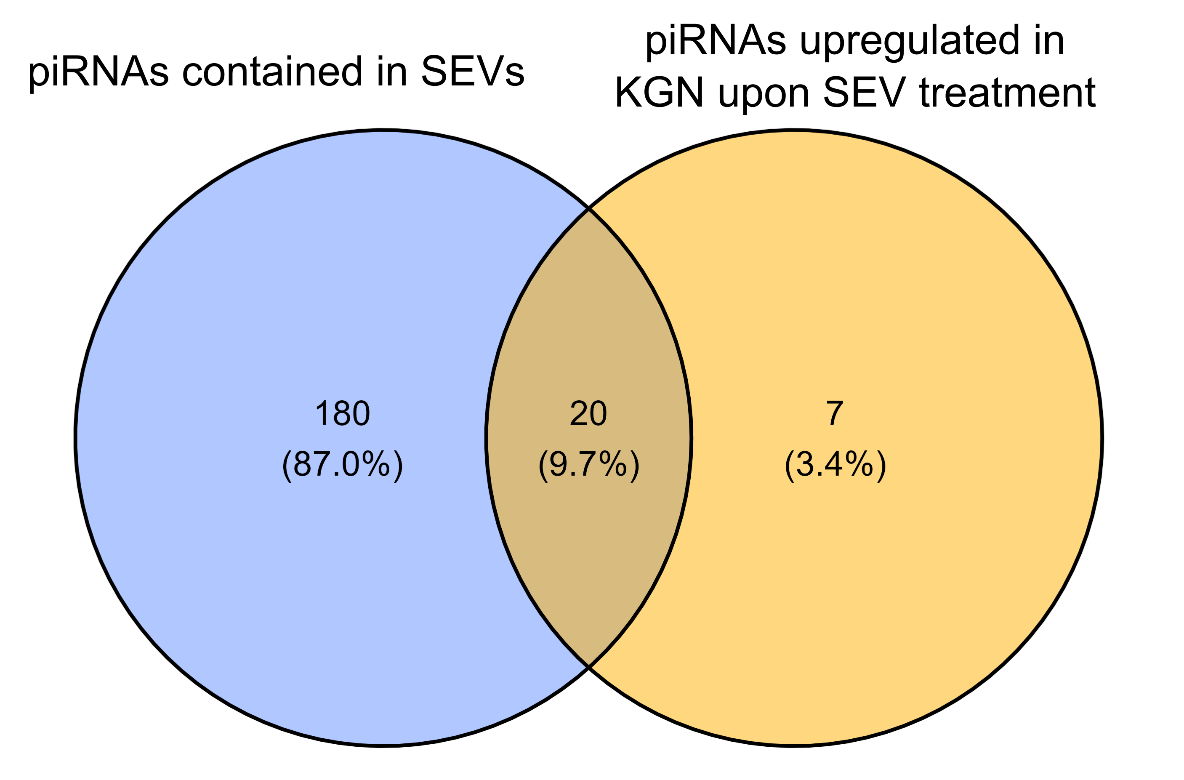

**Supplementary Figure S5.** The overlap between piRNAs upregulated in KGN cells upon SEV treatment and piRNAs contained in SEVs.

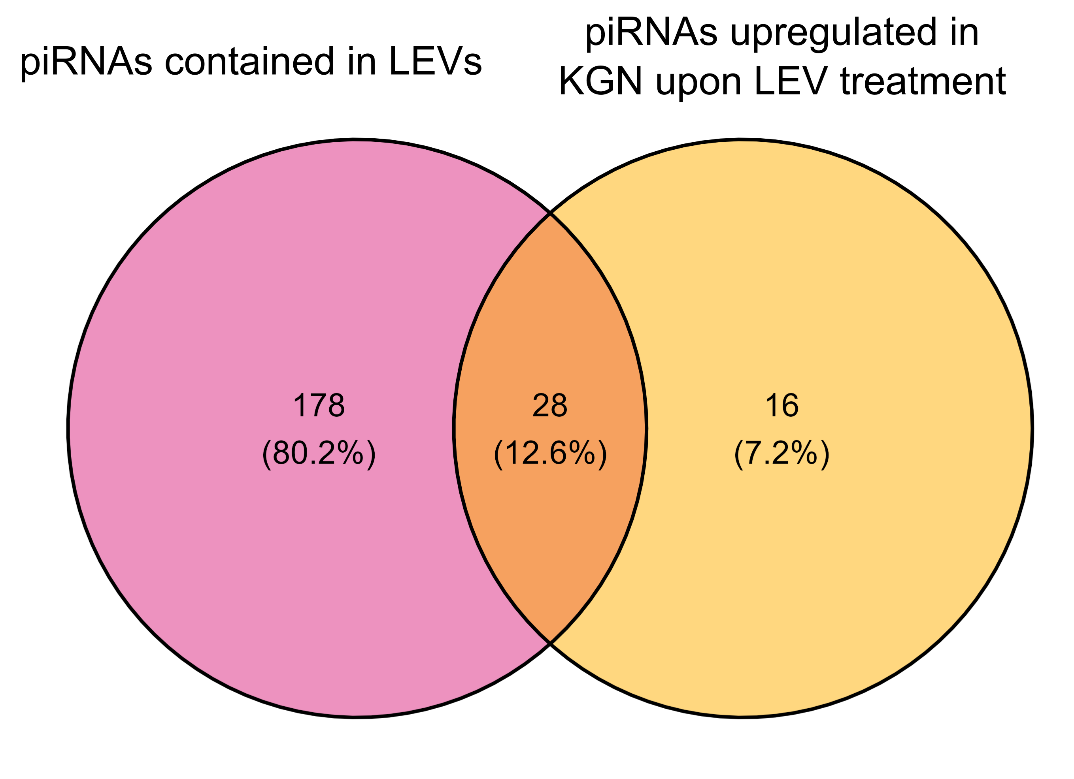

**Supplementary Figure S6.** The overlap between piRNAs upregulated in KGN cells upon LEV treatment and piRNAs contained in LEVs.
